# Integrative, high-resolution analysis of single cell gene expression across experimental conditions with PARAFAC2-RISE

**DOI:** 10.1101/2024.07.29.605698

**Authors:** Andrew Ramirez, Brian T. Orcutt-Jahns, Sean Pascoe, Armaan Abraham, Breanna Remigio, Nathaniel Thomas, Aaron S. Meyer

## Abstract

Effective and scalable exploration and analysis tools are vital for the extraction of insights from large-scale single-cell data. However, current techniques for modeling single-cell studies performed across experimental conditions (e.g., samples, perturbations, or patients) require restrictive assumptions, lack flexibility, or do not adequately deconvolute condition-to-condition variation from cell-to-cell variation. Here, we report that Reduction and Insight in Single-cell Exploration (RISE), an adaptation of the tensor decomposition method PARAFAC2, enables the dimensionality reduction and analysis of single-cell data across conditions. We demonstrate the benefits of RISE across two distinct examples of single-cell RNA-sequencing experiments of peripheral immune cells: pharmacologic drug perturbations and systemic lupus erythematosus (SLE) patient samples. RISE enables straightforward associations of gene variation patterns with specific patients or perturbations, while connecting each coordinated change to single cells without requiring cell type annotations. The theoretical grounding of RISE suggests a unified framework for many single-cell data modeling tasks. Thus, RISE provides an intuitive universal dimensionality reduction approach for multi-sample single-cell studies across diverse biological contexts.

**Highlights:**

- RISE enables tensor-based analysis of single-cell experiments across conditions.
- RISE separates condition-specific effects from cell-to-cell variation.
- RISE provides intuitive isolation of patterns into condition-, cell-, and gene-specific patterns.

## INTRODUCTION

Single-cell measurements have revolutionized our ability to study the variation within and between heterogeneous cell populations^1,2^. As single-cell RNA-sequencing (scRNA-seq) and other high-dimensional single-cell measurement technologies have become increasingly accessible, these technologies have extended to studies including multiple experimental conditions, samples, or subjects. Multi-condition single-cell experiments evaluate how heterogeneous cell populations behave in response to different biological environments, such as how cells respond to drugs and gene knockouts or how cells differ across patients according to disease pathology^3^. There are substantial challenges in the analysis of these data, and appropriate analytical techniques are essential to learning from these studies.

The size and complexity of single-cell data present inherent challenges, including how to deconvolute biological variation within cell populations from variation arising from condition-specific differences. To address these challenges, studies have employed several computational approaches, such as pseudo-bulk techniques, wherein cells are coalesced into population averages, and differential expression is assessed for each population between conditions^4–6^. Other methods enable cell clustering, gene set enrichment, and pseudotime analyses to identify underlying trends in single-cell data^7–10^. Another source of variation includes technical/batch variations from sample processing. A few prominent batch integration techniques have emerged such as Seurat’s canonical correlation analysis using linear combinations of features^11^. Others include Harmony’s iterative soft clustering to align similar cell types across batches, LIGER’s integrative non-negative matrix factorization which identifies shared subspaces between datasets, and deep learning approaches like scVI, a stochastic optimization and deep neural network model^12–14^. However, batch correction methods introduce subjective bias and risk removal of meaningful biological differences or preserving technical noise.

Other methods for modeling such data focus solely on the analytical challenges of identifying biological variation across experimental conditions. One method, PopAlign, leverages a few high-level cell types to align a reference and a test sample with a Gaussian mixture model^15^. A plethora of complementary methods have emerged to detect biological differences between experimental treatments, including PhEMD and HSS-LDA^16–20^. Nevertheless, these methods either make restrictive assumptions about how the data is distributed^15,17,20^; do not directly isolate patterns of variation to specific conditions, cell populations, and genes^16–18^; or reduce the analysis of single-cell variability to a specific number of pre-determined cell types^19^.

Many of these computational methods make use of matrix factorization as a key part of their workflow, as it allows large datasets to be summarized into interpretable and computationally tractable patterns that are each associated with unique sets of cells and genes. However, matrix factorization techniques fail to account for the inherently high-dimensional sources of variation in such studies (variation with regard to cells, genes, conditions) as they require that data be concatenated into a single two-dimensional data matrix where each row corresponds to a cell, regardless of its condition, and the column represents a gene that is measured^21^. When these multidimensional datasets are reduced to two dimensions, the distinction between cell- and condition-associated variations becomes conflated, obscuring the independent contributions of each dimension^22^. This conflation is a fundamental issue for analyzing multi-condition single-cell data and hinders the identification of biologically meaningful patterns.

One approach for analyzing measurements across several dimensions is rearranging the data into a tensor, or a multidimensional array, and using multilinear tensor decomposition techniques^23^. Canonical polyadic decomposition (CPD), one such tensor decomposition technique, approximates a dataset as the sum of the outer product of *n* vectors for an n-dimensional input, with each vector corresponding to a dimension of the data. CPD has been useful in bulk RNA-seq analysis measured across different tissues and patients; where the 3D structure of the data can be approximated by component patterns, each component having three vectors which explain how that pattern is associated with each gene, tissue, and patient, respectively^24^. Where these methods are appropriate, we and others have observed several benefits of a tensor decomposition-based analytical approach^24–26^; tensor decompositions can be more effective at removing noise, isolating distinct variation patterns, and imputing missing values compared to matrix-based techniques^24,27–29^. Most importantly, associating trends in the dataset to specific dimensions helps to interpret the results and therefore derive insights from the underlying experiments^24,30,31^. For example, tensor-cell2cell is a method that associates cell-cell communication signatures with specific experimental samples, using pseudobulk cell type profile information^28^. Dimension-specific associations also make these methods especially effective in data integration, combining datasets with shared experimental parameters^24,32–36^.

Single cell experiments across conditions might be naturally organized into a three-dimensional tensor with axes representing the condition (e.g., samples, perturbations, or patients) in which the cells were collected, individual cells, and each gene measured. However, while conventional tensor decomposition techniques such as CPD assume alignment, or consistent correspondence across dimensions, single-cell technologies do not measure the same cell across conditions. One solution for decomposing data that lacks alignment along one dimension is an approach referred to as PARAFAC2^37^. PARAFAC2 derives a series of projection matrices for each condition to align the data across dimensions, enabling factorization of the data and thus comparative analysis across each sample. Originally developed to analyze data from chromatography experiments, where the elution time of molecules can vary from run to run, PARAFAC2 has been applied in various contexts, including electronic medical health records and spectroscopy experiments^38–40^. In addition, PARAFAC2 is the basis for Fast-Higashi, a computational technique that that addresses chromosomal organization in single cells, specifically addressing the challenge of comparing chromosomes of different sizes^41^.

Here, we demonstrate that our modification of PARAFAC2, Reduction and Insight in Single-cell Exploration (RISE), provides an effective and general solution for integrating multi-condition single cell experiments by dissecting the variation found within and across conditions. RISE improves on the resolution of this analysis by directly modeling individual cells in each condition, enabling the identification of subpopulations of cells with distinct behavior that are missed by existing approaches. RISE is an unsupervised method that does not require external labeling of cells into cell types or categorization of the experimental conditions, thus reducing the number of potentially restrictive assumptions imposed on the data prior to modeling. Both benefits result in enhanced associations with disease status in observational studies, and nuanced insights about the signatures associated with patient features. RISE builds upon the well-developed theoretical foundations of PARAFAC2, with connections to other tensor decomposition algorithms, making RISE readily extensible to other challenges, such as integrating multi-omic or single-cell and bulk measurements. Based on these observations, we propose RISE as a broadly effective solution for analyzing single cell experiments across several samples or conditions.

## RESULTS

### RISE integrates single-cell measurements across conditions through the principle of parallel proportional profiles

Multi-condition single-cell data is typically analyzed by flattening/concatenation, where all single-cell experiments are combined into one large two-dimensional matrix of cells by genes (Fig. 1a). Consequently, once data reduction is performed on this matrix, the association of each component pattern with cells and conditions is confounded. Instead, one might imagine structuring this data as a condition by cell by gene tensor and then applying canonical polyadic decomposition (CPD) to create distinct component associations with each dimension of the single-cell dataset. While CPD is appropriate for data that can be grouped into corresponding cell types across samples (Fig. 1b), single cells exist across in a heterogenous manner across continuum of cell states in gene expression space rather than in discrete, rigid cell types^42^.

**Figure 1.**
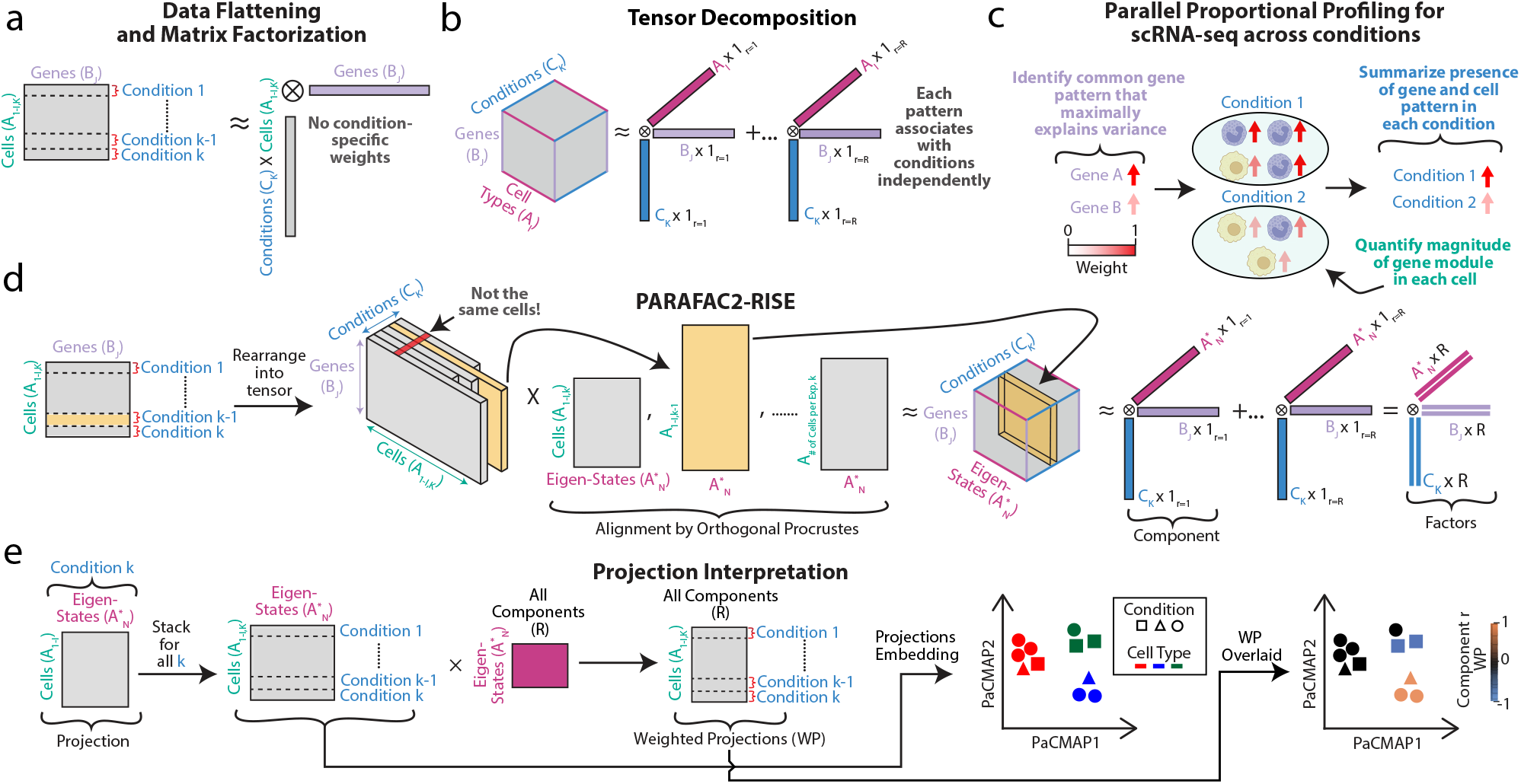
RISE enables the interpretable integration of single-cell measurements across conditions. **(a)** Traditional data reduction methods are applied to flattened multi-condition data in matrix form. **(b)** Instead, tensor decomposition of aligned multi-condition data results in loadings describing the component association with each dimension. **(c)** PARAFAC2 satisfies the principle of parallel proportional profiles which states that latent variable models can be most effectively aligned by identifying common underlying patterns (here, gene signatures) in the dataset that only vary in their proportional magnitude within each condition or sample. **(d)** Multi-condition single-cell experiments can be naturally organized into a staggered three-dimensional tensor; PARAFAC2 uses Orthogonal Procrustes to project cells onto common cell eigen-states and tensor decomposition (CANDECOMP/PARAFAC) to factor this aligned data. **(e)** The projections for each experimental condition multiplied by the eigen-state factors provide an association of each cell and component pattern. The projections alone also summarize the cell-to-cell variation independent of condition-specific effects.

One solution for analyzing such data is Reduction and Insight in Single-cell Exploration (RISE), our extension of PARFAC2 tailored for single cell biology. PARFAC2 builds upon Raymond Cattell’s 1944 principle of parallel proportional profiles, which asserts that patterns across datasets can best be aligned by identifying a shared set of patterns (here, gene signatures) in the dataset that only vary in their proportional magnitude (Fig. 1c)^37,43–45^. In the context of scRNA-seq, this suggests identifying a common set of gene signatures across all cells and conditions and then assigning a magnitude that reflects the strength of that gene signature. This principle is particularly valuable for analysis of scRNA-seq because it allows for the interpretable characterization of cells with subtle variations across experimental samples, even among cells of different cell types. Unlike existing singe-cell integration that forcefully aligns transcriptional profiles between conditions, PARAFAC2 quantifies relative abundances of gene signatures while preserving condition-specific variation to avoid overcorrection of biological differences (Table S1).

PARAFAC2 introduces an additional constraint that enables solutions that are unique beyond trivial scaling and permutation differences, while avoiding restrictive assumptions such as non-negativity or a specific underlying cell distribution^15,43^. Most conservatively, PARAFAC2 assumes similar variable cross-product matrices between experimental conditions, although many successful applications violate this assumption^43^. With scRNA-seq, this translates to the expectation of a roughly similar gene-to-gene covariance structure between conditions^37^. Due to its applications outside the biological sciences, the scaling and convergence performance of PARAFAC2 has already been widely explored alongside additional constraints such as restriction to only non-negative factors^37,43,46–49^. We have developed RISE by optimizing PARAFAC2 specifically for scRNA-seq data by including refined parameter initialization, strategic parameter selection, use of Nesterov acceleration, and post-factor modifications (see in Methods).

As part of its fitting process, PARFAC2 defines a standardized reference frame across conditions. This is accomplished by Orthogonal Procrustes, an algorithm that uses a linear projection to summarize data into a predefined number of cell identities, here termed “eigen-states” (Fig. 1d). This results in one projection matrix per sample, which converts all the cells in that sample into the common eigen-states. Thus, an eigen-state represents a weighted collection of cells with a shared set of gene signatures. This identifies cells which share expression profiles present across experimental conditions. The projections only contain information about cell identity, not condition-to-condition variation; they represent which cells are relevant when encompassing an eigen-state, thereby mapping cells to an underlying common latent space. Thus, the projections provide a common basis to compare cells with similar or dissimilar gene expression behavior across all samples.

After projecting the data into an aligned tensor, the projected tensor is factored via CPD^23^. CPD applied to the eigen-state intermediate tensor results in the weighted sum of rank-one vector outer products for the genes, eigen-states, and conditions. Similarly to PCA, each component, or a set of three vectors, describes a specific variation pattern in the data. Each component is defined by three separate vectors defining the association of variation to specific genes, eigen-states and conditions. The combination of all vectors across one dimension are the factors and define how components vary across that one dimension. Because both the cell alignment and factorization are initially unknown, the two steps of cell alignment via the projections and CPD factorization are iteratively repeated until the solution converges (Fig. S1, Methods). Importantly, the CPD factorization enforces a low-rank structure, and the projection matrices map each experimental condition onto this shared low-dimensional space.

The PARFAC2 algorithm yields projections for each experimental condition and three factor matrices for the genes, eigen-states, and conditions. As part of RISE, we adjusted the scaling, sign interpretation, and order of the components and eigen-states to improve factor interpretation of the PARAFAC2 model (see in Methods). To determine which cells are highly associated with a specific component, the eigen-state factor matrix can be converted into a cell-specific factor matrix for each component (Fig. 1e). These “weighted projections” are derived by multiplying the eigen-state factor matrix and the projections across all conditions. We can then perform non-linear dimensionality reduction to visualize the projections in a low-dimensional embedding (Fig. 1e). This projection latent space describes cell-to-cell variation, separating any condition-specific effects into the factors. To illustrate which cells are associated with a given component, we can overlay the weighted projections on the projection embedding. Collectively, this enables RISE to effectively identify biological variation across experimental conditions and cellular states common across all conditions.

### RISE separates cell-to-cell from condition-specific variation

To explore the utility of RISE, we first examined scRNA-seq data from Chen et al., a study wherein human peripheral blood mononuclear cells (hPBMCs) from one healthy donor were treated with a panel of 40 drugs from an FDA-approved immunomodulatory compound library, along with 6 control conditions (Fig. 2a)^15^. Previous analysis, using mixture modeling, isolated the transcriptional changes induced by each drug within two broad immune cell populations: T cells and monocytes (MO). Compounds were ranked based on their dissimilarity to the unperturbed controls to determine which experimental conditions most altered gene behavior. We surmised that RISE might improve on the resolution of the previous analysis by revealing smaller subpopulations with unique gene expression responses, as well as shared or distinct responses across drug perturbations. Because PCA represents the optimal linear dimensionality reduction technique and a baseline for variance explained, RISE was compared to PCA. RISE and PCA explained similar amounts of variance using the same number of components, despite RISE being a more mathematically constrained model (Fig. S2a)^37^.

**Figure 2.**
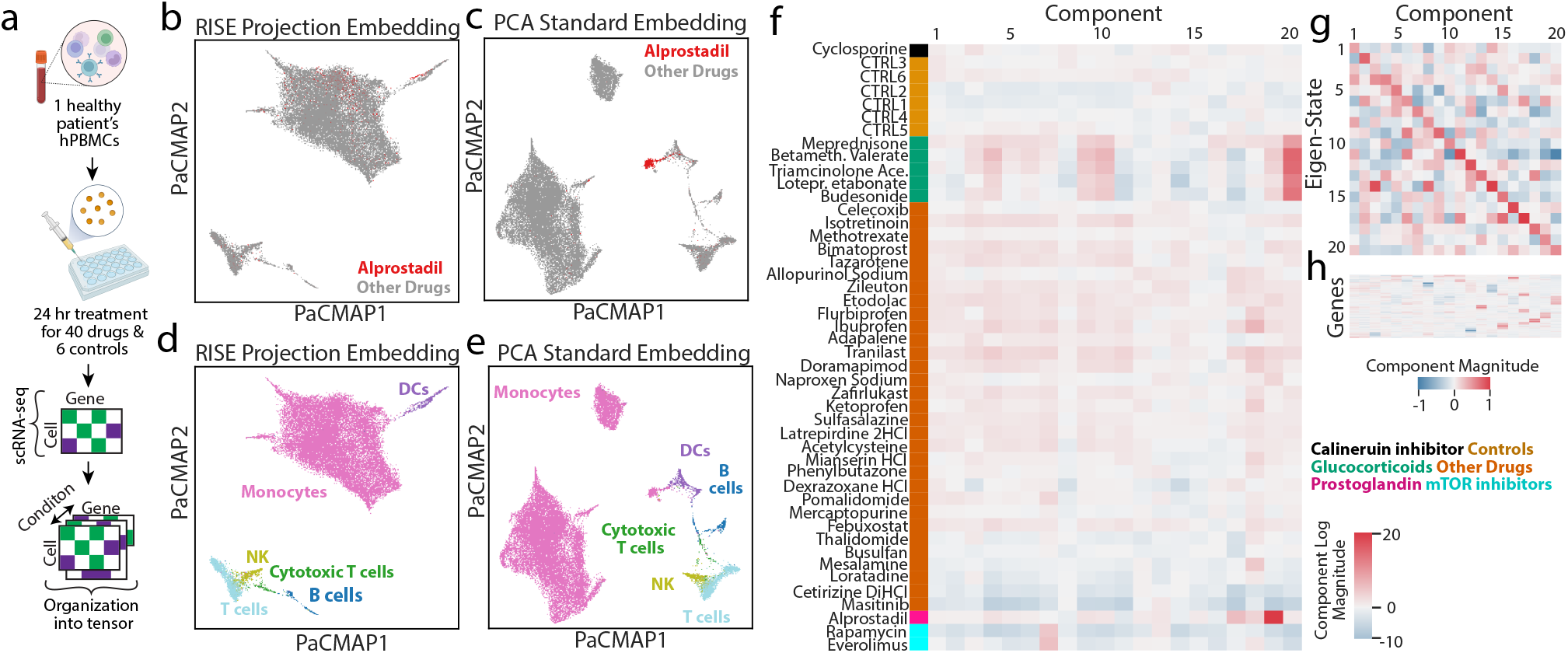
RISE defines cell-to-cell and drug-specific variation in a drug perturbation scRNA-seq experiment. **(a)** hPBMCs were treated with 40 FDA-approved drugs, including glucocorticoids, prostaglandins, mTOR inhibitors, and calcineurin inhibitors, alongside six controls. **(b–c)** PaCMAP embedding of the (b) RISE projections and (c) PCA-based dimensionality reduction, with cells from the alprostadil treatment condition highlighted. **(d–e)** PaCMAP of the (d) RISE projections and (e) PCA-based dimensionality reduction, with cells colored by their cell type annotations. **(f–g)** The (f) drug condition, (g) eigen-state, and (h) gene factors for each component. The genes factor matrix is filtered for genes having an effect magnitude of 0.8 or more within one of the components. The conditions factors are non-negative and have been log transformed to better show weights relative to the control conditions.

After fitting RISE to the Chen et al. data with 20 components, we visualized the cell-to-cell variation using a PaCMAP embedding of the RISE projections (Figs. 2b, d)^50^. We used PaCMAP, a non-linear embedding method, for cell embedding, as it has been shown to more reproducibly preserve local relationships and prevent distortion of global structures as compared to other methods like t-SNE and UMAP. As a basis of comparison, we also plotted the PaCMAP embedding of the PCA scores with the same number of components (Figs. 2c, e). Whereas the PCA-based embedding resulted in clusters defined by their drug perturbation, the RISE projections-based embeddings distributed cells from each condition across the embedding space (Fig. 2b, 2c). Similar results were observed for other compounds, suggesting that the projection matrices generated using RISE represent patterns that have been deconvolved from condition-specific varying, unlike the factors generated by PCA (Fig. S2b–e). We clarify that RISE is not a batch correction method, but rather an approach that maps cell-to-cell variation in the projections and defines condition-specific differences in the underlying factors, as supported by the above results. Manually annotating the cell types revealed that the RISE projection embeddings resolved many cell subpopulations despite condition-to-condition differences (Fig. 2d, e). To quantify these effects, we observed that the PCA standard embedding resulted in poorer integration accuracy in both batch integration and conservation of biological variance in immune cell types, most likely due to clustering cells based on perturbation-specific gene expression responses (Fig. S2f)^51^. As expected, given this experiment involves short-term treatment of PBMCs from one donor, abundant cell types were similarly represented across all conditions (Fig. S3a-g).

Like PCA, RISE results in a series of component patterns, but with vector loadings representing the association of the pattern along the three dimensions of the data: drug perturbations (Fig. 2f), eigen-states (Fig. 2g), and genes (Fig. 2h). Each component pattern can be traced across the dimensions in the dataset and illustrates a distinct biological trend. A small fraction of genes was highly associated (i.e., highly weighted) with each component, consistent with our expectation that the effects of immunologic drugs on various cell types are defined through specific gene signatures (Fig. 2h).

To determine whether RISE condition and gene factors trends were robust to changes in cell number, we applied RISE to subsets of the drug perturbation panel. Using the Factor Match Score (FMS) to quantitatively measure the similarity of the factor matrices, we found that the results were essentially unchanged with the dataset reduced to as much as half its original size (Fig. S4a)^49^. This highlights the fact that the RISE eigen-states indeed correspond to robust signatures in the underlying data, as they are reproducible when only a representatively sampled subset of the cells are present. Using a bootstrapping strategy, we similarly observed consistent factor results at varying number of components, though a decreasing trend in the FMS at the highest number of components, as expected (Fig. S4b). These results indicated that RISE provides reproducible results and that we were not overfitting these data. Having confirmed the computational suitability of RISE for modeling our data, we next wondered what biological insights RISE can identify.

### RISE identifies cell-to-cell, condition-specific, and context-dependent gene expression variation

Having assessed the reliability of the RISE factorization, we next inspected the patterns of gene expression variation that were identified (Fig. 2f–h). To interpret the biological significance of a component, one can inspect the associations across specific conditions, genes, and cells. Cell types were manually annotated using cell type marker genes at higher resolution to improve interpretation of the RISE components (Fig. 3a, Methods). One of the first advantages of RISE is the ability for components to identify common cells with similar gene expression profiles across experimental conditions. For instance, we plotted the component 15 weighted projections scores of individual cells on the PaCMAP embedding and found that plasmacytoid dendritic cells (pDCs) were selectively and highly weighted (Fig. 3e). FXYD2, SERPINF1, and RARRES2 are genes which have been identified as selectively expressed in pDCs in the literature and Human Protein Atlas^52–54^. These three genes were identified as the most strongly weighted in component 15, and further examination of these genes confirmed their specificity in pDCs (Fig. 3b). Examining the conditions factor, component 15 was largely invariant in its association with drugs, suggesting this component represents a gene expression module that is equally present across all conditions (Fig. 3c). Taking these findings together, we can conclude that component 15 represents the presence and unique gene expression of pDCs across all drug treatment conditions. Only 5 pDCs were found within each condition on average, demonstrating RISE’s sensitivity to low-abundance cell populations, as well as its ability to reveal meaningful variation outside of contrasts between cases and controls (Fig. S5m). In addition to pDCs, RISE identified that component 12 was strongly associated with the amount of B cells in each drug perturbation experiment based on the genes identified by this component, including MS4A1A and CD79A (Fig. S6l, S7a, Table S2a). To simulate data where a cell population is differentially abundant in an experimental condition, we synthetically reduced or removed B cells from a drug perturbation. The resulting RISE factorization showed a reduced weight for component 10 in the altered condition, demonstrating how RISE identifies experimental conditions with differentially abundant cell populations within a condition (Fig. S7b, c).

**Figure 3.**
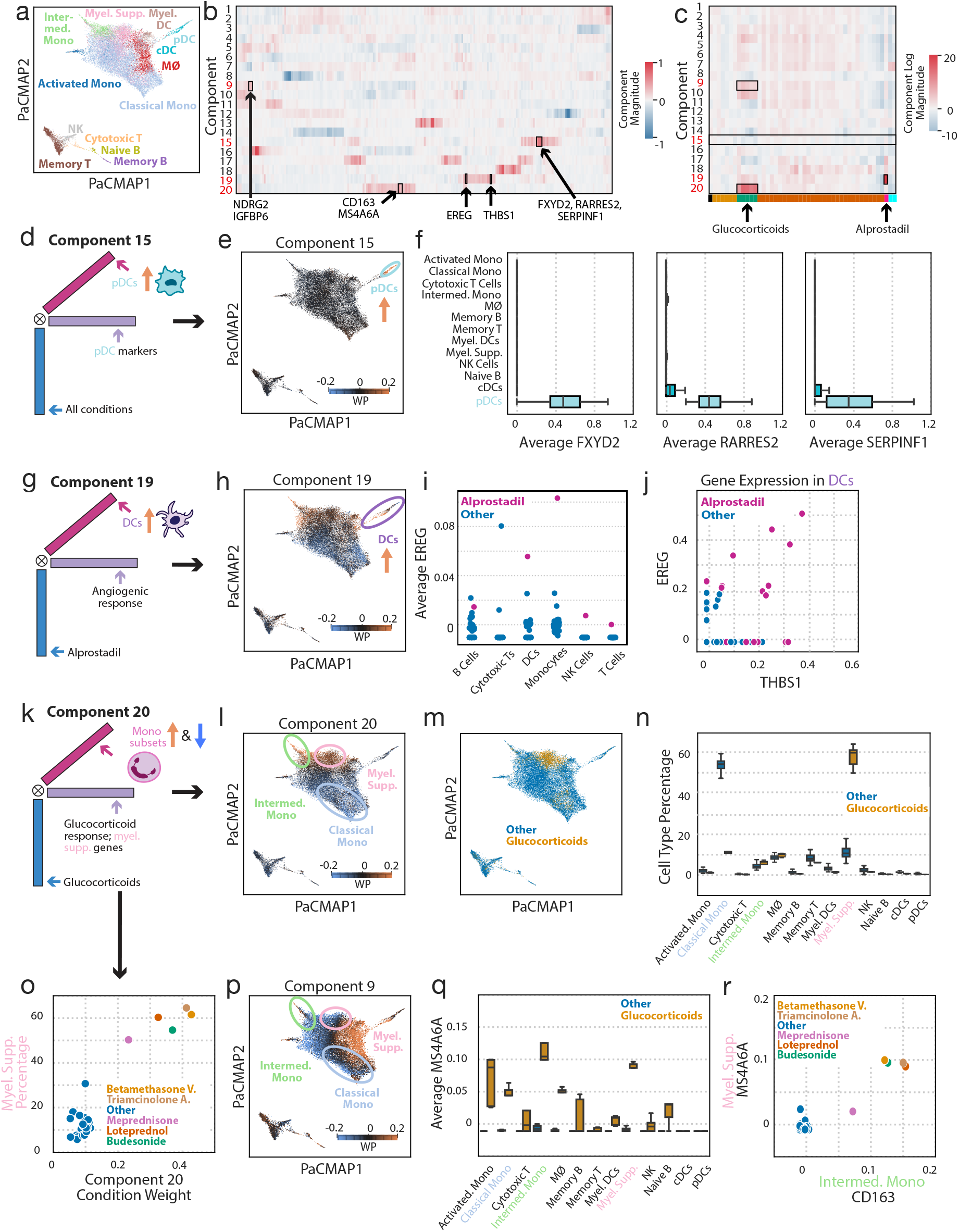
RISE analysis of drug perturbations reveals both broad and highly localized gene expression responses. **(a)** PaCMAP of the RISE projections, with cells colored by their high-resolution cell type annotations. **(b, c)** The (b) gene and (c) condition factors matrices with select genes and conditions highlighted. **(d, g, k)** Schematics of the biological significance of components (d) 15, (g) 19, and (k) 20 describing each component’s association with genes, eigen-states and conditions. **(e, h, l, p)** PaCMAP of the RISE projections, with cells colored by the weighted projections (WP) for components (e) 15, (h) 19, (l) 20, and (p) 9. **(f)** Per-condition average expression of FXYD2, RARRES2 and SERPINF1, stratified by cell type. **(i)** Average gene expression of EREG across drug conditions for low-resolution cell types. **(j)** Expression of THBS1 and EREG in individual dendritic cells, with alprostadil-treated cells labeled. **(m)** PaCMAP of all cells colored by whether they were treated with glucocorticoids. **(n)** Proportion of cells classified as each cell type for each condition, stratified by glucocorticoids versus all other drugs. **(o)** Percentage of cells classified as classical myeloid suppressors for each condition and their corresponding component 20 weight. **(q)** Average expression of MS4A6A within each cell type stratified by whether cells were treated with glucocorticoids or all other drugs. **(r)** Average CD163 expression in intermediate monocytes versus average MS4A6A expression in myeloid suppressors for each condition.

In addition to representing the abundance of specific cell populations, RISE also identified patterns associated with specific drug perturbations. For example, component 19 was strongly and uniquely associated with alprostadil, a prostaglandin inducer of vasodilation and angiogenesis (Fig. 3c) ^55^. Component 19 most strongly weighted the DC population (Fig. 3h), and the component-associated genes included various ones relating to angiogenesis such as THBS1, VMO1, CXCL5, EREG and VEGFA (Table S2b). VEGFA and EREG have been previously confirmed to be upregulated in response to prostaglandin stimulation^56,57^. Alprostadil-treated DCs expressed the most EREG and THBS1, and were the only DCs to co-express these, demonstrating RISE’s ability to identify coincident patterns of gene expression within individual populations (Fig. 3i, 3j). Thus, component 19 represents an angiogenic gene module expressed in DCs that is uniquely induced by alprostadil and illustrates RISE’s ability to model condition- and cell population-specific expression patterns.

RISE’s tensor-based structure can also identify expression modules shared across several drug perturbations. For example, component 20 has a strong relationship with all glucocorticoids in the dataset. Glucocorticoids are steroid hormones commonly used to treat inflammation, autoimmune diseases, and cancer and are known to induce broad transcriptomic effects across immune cell populations^15^. The variance encoded by component 20 is strongly represented in the monocyte population, with distinct effects by monocyte subtype (Fig. 3l). Here, intermediate MO and myeloid suppressors had positive weights, while classical monocytes (MOs) had negative weights, suggesting that the gene expression and abundance information encoded by component 20 is divergent across these subpopulations. Cells in the myeloid suppressor cluster were more abundant in the glucocorticoid-based experiments, mirroring the cell specificity of component 20 (Fig. 3m). Indeed, when examining the cellular composition of samples treated with glucocorticoids, we observed a substantial decrease in MOs, and an increase in myeloid suppressor cells (Fig. 3n). The distributions of all other cell types remained consistent across conditions, suggesting that our projections-based embedding was not simply being informed by drug perturbation, as the PCA-based embedding was. The component 20 weight positively correlated with the proportion of cells labeled as myeloid suppressors (Fig. 3o). This suggests that component 20 captures glucocorticoid-induced expansion of immunosuppressive phenotypes within the myeloid compartment^58–60^.

Although component 9 was also glucocorticoid-specific, it isolated distinct gene signatures and cell associations as compared to component 20 (Fig. 3c/p, Table S2a, S2b). Component 20 described glucocorticoid-induced CD163 and MS4A6A expression across MOs and other cell types, as seen in earlier studies (Fig. 3b, q, S7) ^61^. CD163 upregulation in intermediate MOs correlated with MS4A6A expression in myeloid suppressor cells (Fig. 3r), suggesting coordination between these cell types and demonstrating RISE’s ability to identify shared expression changes between populations. These observations indicated that RISE could identify both condition-specific expression patterns and cellular abundance differences, as demonstrated by its myeloid subtype analysis. However, because RISE simply identifies gene expression signatures that vary in abundance, each component must be examined to determine whether it represents cellular abundance of a cell type or cellular expression differences. Overall, RISE was able to effectively separate distinct and highly interpretable patterns of gene expression variation, whether cell type-specific (Fig. 3e/f), condition- and cell type-specific (Fig. 3h/i), or even conditions with shared alterations among subpopulations of cells (Fig. 3o–q). Here, we only highlight a select few patterns; we also found that the other components identified by RISE identify relevant biological insights as well, each of which is explored in Table S3.

### RISE improves the resolution and accuracy of transcriptomic associations with systemic lupus erythematosus

With an understanding of the benefit RISE can provide within perturbational studies, we next examined its potential to separate cell-to-cell from patient-to-patient variation and to generate interpretable patterns which are predictive of patient status within an observational cohort study. To do so, we focused on a recent study that characterized hPBMCs from a cohort of 162 systemic lupus erythematosus (SLE) patients and 99 healthy controls by scRNA-seq^62^. Multiple scRNA-seq samples were collected from some patients, resulting in 209 SLE and 145 healthy samples. Samples varied widely in overall cell number (Fig. S8a). The previous analysis aggregated single-cell gene expression into pseudobulk profiles for high-resolution cell types and performed differential expression analysis. Approximately 300 differentially expressed genes for groups of immune cells (e.g., all myeloid cells or all immune cells) were used as features for a model distinguishing SLE from healthy samples^62^.

We first investigated RISE’s general explanatory and associative power. RISE explained an increasing fraction of the dataset variance with more components (Fig. S8b). To select an appropriate number of components, we used a logistic regression classifier to associate the patient (i.e., condition) factors with SLE status. Cross-validation was performed identically to the previous study, using batch-specific training and testing; briefly, we trained the logistic regression model on one of the four processing batches of data and then used the patient sample factors to predict the status of the left-out batches^62^. Increasing the component number improved training accuracy for SLE up to 30 components (Fig. 4a, S8c). Bootstrapping the samples or using leave-one-batch-out cross-validation did not strongly alter these observations (Fig. S8d-f). A 30-component RISE model was predictive of SLE status using the same cross validation as the previous authors and resulted in an AUC ROC of 0.94 (Fig. 4b). This approach outperformed both the previously used pseudobulk prediction analysis, and our analysis using pseudobulk gene expression and cell type composition as predictors to classify SLE status. The 30 components were variously associated with the patients’ samples, eigen-states, and genes (Fig. S8g). Examining the cell-to-cell variation within the projections via the PaCMAP embedding, we observed effective separation of the expected immune cell types in hPBMCs, both at lower and higher resolution (Fig. S8h, 4c). In addition, we compared RISE’s scalability and computational efficiency and found that it compares favorably to other single-cell analysis tools (Fig. S9a-d).

**Figure 4.**
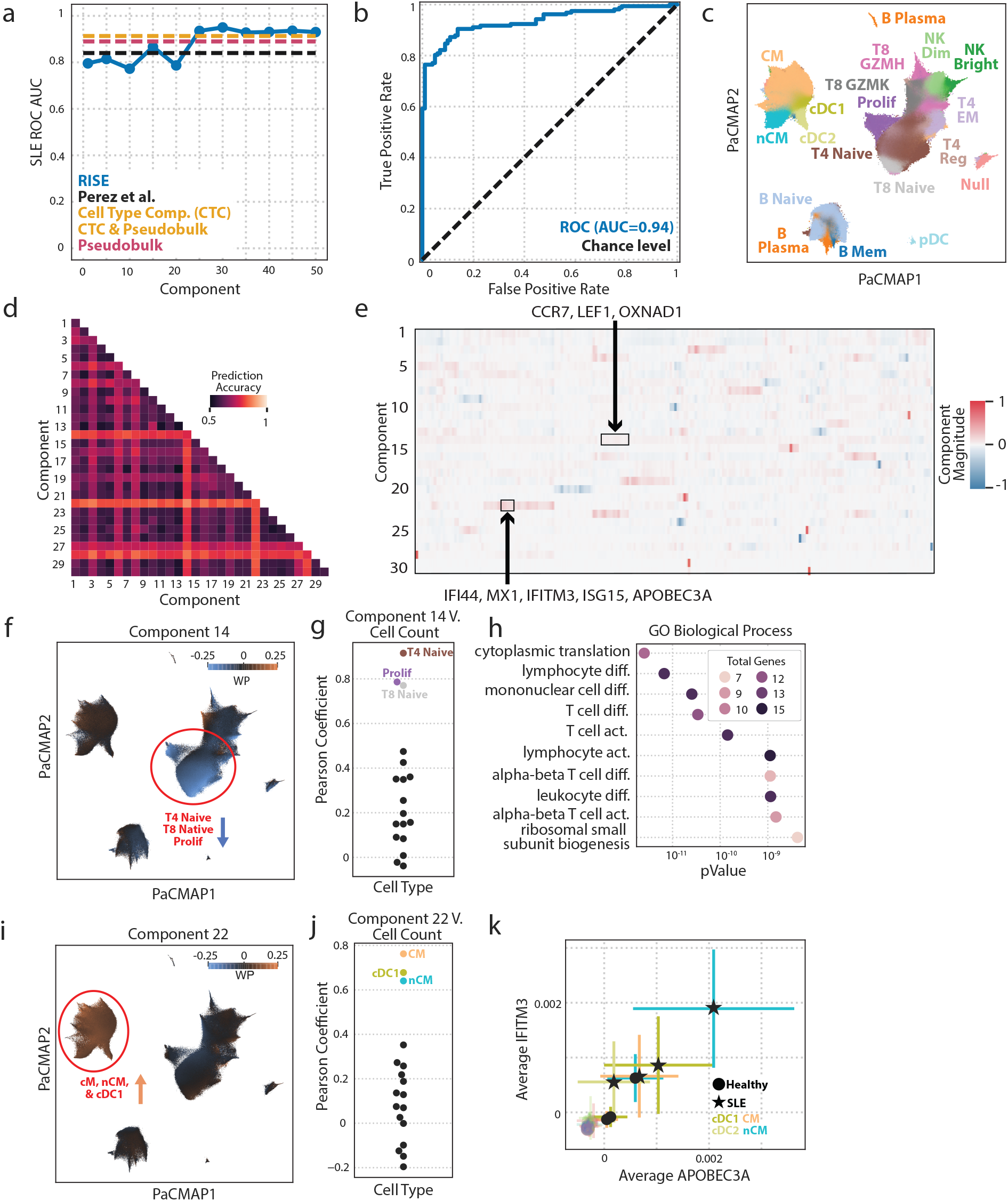
RISE improves the resolution and accuracy of transcriptomic associations with systemic lupus erythematosus. **(a)** ROC AUC of a logistic regression classifier, using the patient factors, predicting SLE status. RISE rank was varied between each model. **(b)** ROC curve of a logistic regression classifier, using the patient factors, predicting SLE status with a 30-component RISE model. Cross-validation was performed by leave-one-processing-batch out. **(c)** PaCMAP of the RISE projections, with cells colored by their higher resolution cell type annotations. **(d)** SLE prediction accuracy of an unpenalized logistic regression model trained on pairs of components. **(e)** The gene factors matrix using 30 components, with genes from components 14 and 22 highlighted. **(f, i)** PaCMAP of the RISE projections, with cells colored by the weighted projections of components (f) 14 and (i) 22. **(g, j)** Pearson correlation coefficient based on the relationship of cell type count per patient against corresponding patient sample weightings of components (g) 14 and (i) 22. **(h)** Gene set enrichment analysis (GSEA) biological process results for component 14 gene module (top 30 positively weighted genes), where dot color represents the number of genes overlapping with the gene ontology. **(k)** Average expression of IFITM3 and APOBEC3A for low-resolution cell types in healthy and SLE samples.

An L1-penalized logistic regression model identified components relevant for differentiating SLE sample status (Fig. S9f). However, L1 regularization eliminated components that could possibly differentiate SLE status. We therefore also assessed the classification accuracy using all pairwise combinations of components (Fig. 4d). Notably, components 14, 22, 27, and 28 led to accurate classification in reduced models. We again traced the components across the cell and gene dimensions to interpret the significance of each RISE pattern (Fig. 4e, Table S4a, b).

We first investigated component 14 as it was most strongly SLE-associated. Component 14 selectively associated with Naïve CD4+ and CD8+ T cells, as well as a subcategory of proliferating T cells (Prolif) (Fig. 4f). The sample associations of this component directly correlated with the abundance of these annotated lymphocyte populations (Fig. 4g). Consistent with explaining the abundance of these cell populations, the genes associated with component 14, including CCR7, LEF1, and PRKCQ-AS1 (Fig. 4e), are crucial for early T cell development, as seen similarly by the previous analysis of the data^63,64^. Gene set enrichment analysis (GSEA) of the 30 most weighted genes for component 14 confirmed an association with T cell activation and lymphocyte differentiation (Fig. 4h, Table S4b).

Component 22 was also associated with SLE status and corresponded to the myeloid populations of CM, nCM and cDC1 (Fig. 4i). Similarly to the T cell development pattern, component 22 correlated with the cell abundance of these specific myeloid populations (Fig. 4j). This component was associated with interferon (IFN)-induced genes, including IFI44, MX1, IFITM3, APOBEC3A, and ISG15 (Fig, 4e). Other components overlap in these IFN genes, such as component 6, but highly weight distinct cell populations (Table S4a, b). The IFN-stimulated cell types identified by component 22 uniquely express IFITM3 and APOBEC3A in combination (Fig. 4k). Thus, RISE recovered several known alterations associated with SLE^62,65^.

### RISE identifies novel single-cell patterns associated with SLE

We next explored whether other components might yield novel insights into SLE-associated mechanisms (Fig. 5a). We first inspected component 4 due to its association across several cell types, including GZMH+ CD8+ T cells and several NK populations (Fig. 5b). SPON2, FGFBP2, GZMB, PRF1 and GZMH were among the highly weighted genes associated with component 4 (Fig. 5a, S11). Many of the genes identified by this pattern also overlap with the genes previously used to define a cytotoxic signature for GZMH+ CD8+ T cells^62^. Using the same cytotoxic signature to quantify this gene module as the original authors, we found that other immune cell types also expressed this signature, including NK Bright, NK Dim, and T8 GZMK+ CD8+ T cells, as identified by component 4 (Fig. 5c). This finding demonstrated that the RISE can uncover gene expression changes that are coordinated between cell populations which would have been missed in per-population analyses^66,67^.

**Figure 5.**
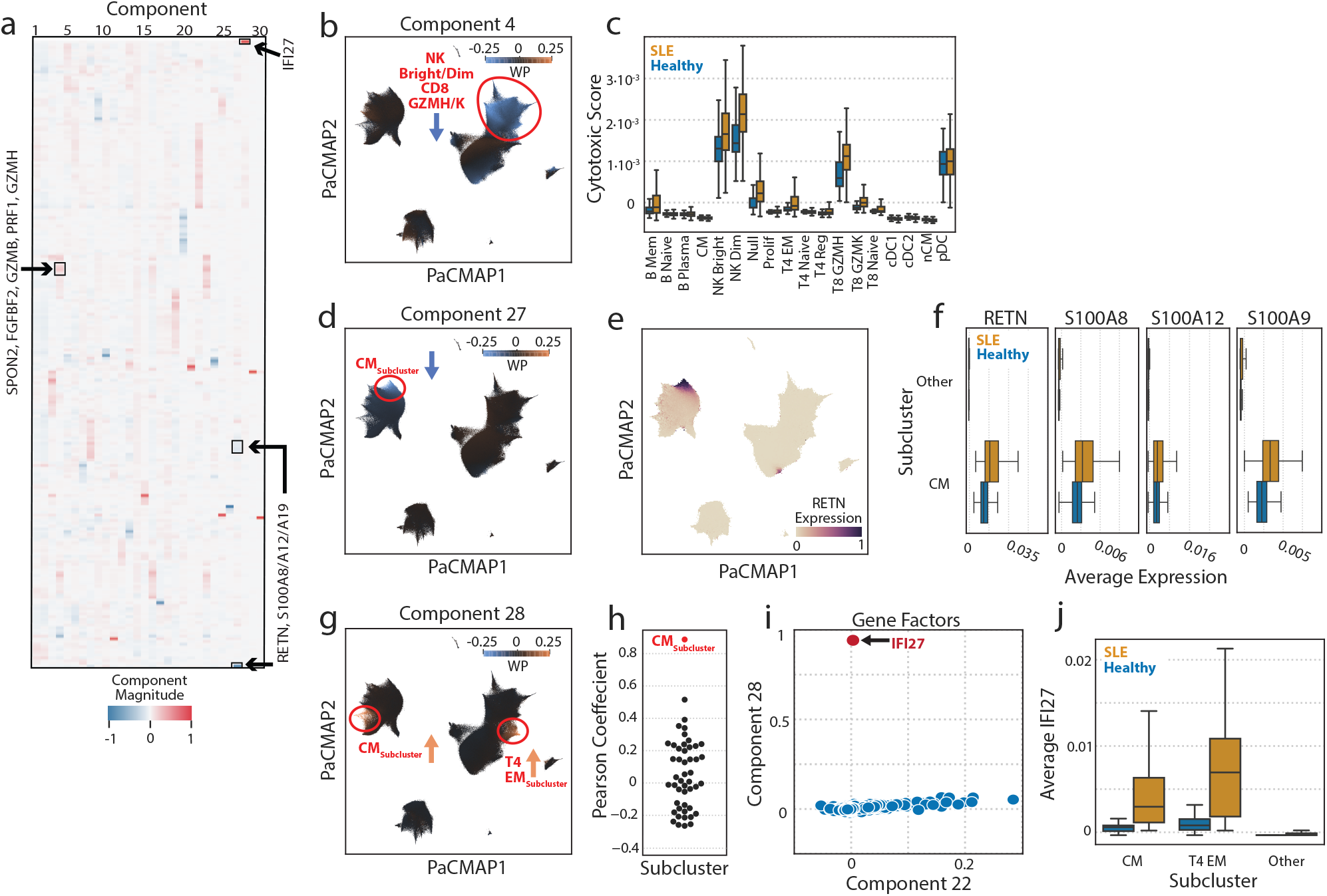
RISE identifies high-resolution subpopulation and gene signatures associated with SLE. **(a)** The genes factors matrix with select genes highlighted. **(b, d, g)** PaCMAP of the RISE projections, with cells colored by the weighted projections of components (b) 4, (d) 27, and (g) 28. **(c)** Cytotoxic gene score stratified by each cell type. **(d)** Normalized RETN expression of single cells overlayed on PaCMAP embedding **(f, j)** Average (g) RETN, S100A8, S100A12, S100A9, and (k) IFI27 gene expression classified by SLE status and subcluster. **(h)** Pearson correlation coefficient based on the relationship of subcluster cell count and condition sample weightings per patient for component 28. **(i)** Correlation between component 22 and 28 gene weightings, with IFI27 highlighted.

In addition to identifying coordinated changes across cell populations, we found that RISE identified gene modules in subpopulations of cells that would be missed by a pseudobulk approach. For example, component 27 was highly SLE associated and specifically weighted a small subcluster within the CM population comprising 1% of the total population of cells per patient sample (Fig. 5d). The most highly weighted gene in this component, RETN, was found to be uniquely expressed in cells identified by high component 27 weights (Fig. 5a, 5d, 5f). In addition to RETN, the genes S100A8, S100A12, and S100A9 were highly weighted, which others have identified as relevant to IL-6 signaling and activation of toll-like receptors 4 and 7 in SLE patients, prolonging inflammation (Fig. 5a, S11) ^68–71^. The component 27-high CM subpopulation is distinct in its shared expression of RETN, S100A8, S100A12, and S100A9 as compared to all other cells (Fig. 5g). RISE revealed the increase in gene expression of the inflammatory module within a subset of classical monocytes, a nuance which would been obscured in a pseudobulk analysis that examined all myeloid cells.

Furthermore, RISE can identify subpopulations of traditional cell types between cell types with similar expression signatures. Notably, component 28, which also had high association with SLE status (Fig 4d), is highly specific to CM and CD4+ effector memory T cell (TEM) subpopulations (Fig. 5h). The component 28 sample weightings were most correlated with the cell abundance of the CM subpopulation and component 28 identified many IFN genes including IFI27, IFI6, ISG15, IFITM3 and IFI44L (Fig 5c). Although some of these genes are highly weighted in other components like in component 22, component 28 strongly weights IFI27 which is specific to these CM and CD4+ TEM subpopulations (Fig. 5i, S11). As expected, the subclusters for both CM and CD4+ EM have a higher average expression of IFI27 in SLE patient samples (Fig. 5j). Thus, one can conclude there are subclusters of both CM and CD4+ effector memory T cells that share this unique expression of IFI27, along with other type 1 interferon regulated genes. Indeed, IFI27 has been suggested as a potential biomarker for SLE in CD4+ T cells and monocytes^65,72–74^. However, our finding hints that not all cells categorized as CD4+ EM and CM contribute to this unique expression gene module.

## DISCUSSION

Here, we demonstrated that RISE improves the investigation and understanding of multi-condition single cell experiments. By aligning cells between conditions during the model fitting process, single-cell data can be directly factored using tensor decomposition, thus preserving the experimental structure more faithfully than matrix factorization methods would. Because RISE builds upon PARAFAC2, an unsupervised data reduction method, no prior information is required about cell identity or the types of experimental conditions, making RISE unbiased as compared to other models that utilize a reference atlas^75,76^. In both the Chen et al. and Perez et al. studies above, RISE identified transcriptional patterns that were shared across conditions as well as those that were specific to a subset of conditions. These patterns of expression were quantitatively linked to individual cells making them highly interpretable. Overall, these results show how a tensor-based approach to single-cell data leads to unique insights that cannot be identified by dimensionality reduction in matrix form or standard pseudobulk analyses, both which obscure biological relationships across conditions. RISE eliminates the need to define the difference between technical and biological variation by characterizing patterns of variation across single cells and experimental conditions in a data-driven manner. RISE thus serves as an alternative single-cell analysis tool that effectively isolates and deconvolutes variation arising from cells, genes and experimental samples, without the need to perform batch correction, or to organize data in a way that obfuscates patterns across dimensions (Fig. 6).

**Figure 6.**
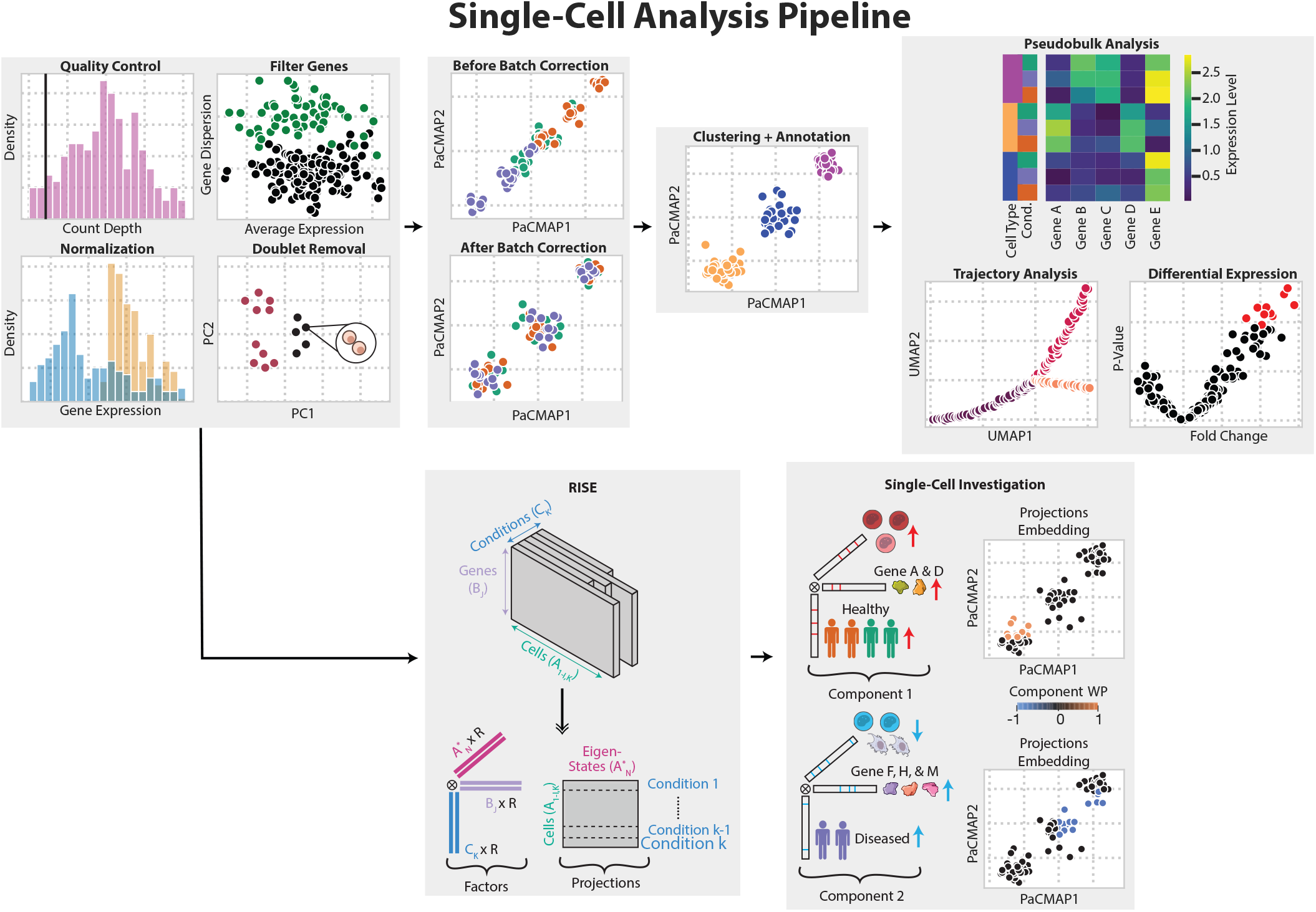
RISE presents an innovative, distinct single-cell analysis method to map interconnected variance in gene modules and cellular populations across experimental samples, providing a comprehensive framework for understanding complex cellular heterogeneity.

The advantage of analyzing single cell experiments across conditions with RISE was first demonstrated for the exploratory analysis of drug perturbations. While earlier analysis demonstrated the ability to identify perturbed populations at a high level, distinguishing T cells and monocytes, the RISE components provide quantitative information about perturbations integratively and at high resolution. Despite not requiring cell annotations, RISE isolated rare subpopulations, such as pDCs, and their canonical markers including FXYD2 and RARRES2/chemerin^77^. It also distinguished several qualitatively distinct glucocorticoid responses, including conversion of cell identity to myeloid derived suppressor cells and variation in the types of responses across monocyte subsets^78^. While prior analysis localized Alprostadil effects to monocytes overall, RISE demonstrated that DCs were most affected by Alprostadil treatment and identified an associated pro-angiogenesis expression signature.

The strengths of RISE were especially evident in more complex single-cell transcriptomic data from SLE patients. While earlier analysis of this data relied on pseudobulk methods, we demonstrated that by directly modeling the single cell data, we were able to improve our identification of patterns which were predictive of SLE status. RISE confirmed known alterations in naïve T cell and monocyte abundances, while also recovering differences in CD8+ T cell maturation, proliferating T cells, and cDC1 cell abundances described by others^67,79–81^. Our analysis also uncovered a coordinated cytotoxic and exhaustion-like GZMH+ expression program that extends beyond CD8+ T cells to NK cells, B cells, and CD4+ T effector memory cells, the last of which have been recently found to be associated with SLE disease progression^66,82^. Additionally, RISE highlighted gene expression modules in a small subset of monocytes with unique expression of RETN and S100A genes. Other studies have suggested S100A8 and S100A9 as potential biomarkers for SLE and this analysis points to subpopulations of CM as responsible for this observed pro-inflammatory behavior in SLE^68,69^. Similarly, RISE identified upregulated coordinated changes of IFI27, another putative SLE biomarker, in specific subsets of monocytes and CD4+ T effector memory cells^83,84^. Many SLE studies have shown the autoimmune disease is characterized overall by a type 1 IFN-regulated response, but RISE delineated distinct IFN signatures according to their gene patterns and cellular localization^85,86^.

Despite the popularity of pseudobulk analysis, these results demonstrate several of the limitations of this approach. One must choose a level of detail at which to annotate the cells, with the ultimate results and even basic features such as statistical power affected by the number of populations the cells are split into. These results are reliant on accurate cell annotations, and small fractions of inconsistently annotated cells could lead to artifacts in differential gene expression analysis. Clustering algorithms are often employed to form populations, which are then annotated. However, there are no guarantees that clustering populations align with their biologically significant counterparts, especially in the presence of technical artifacts between batches of cells^87^. Furthermore, particularly in smaller studies, rare cell-types are often unidentifiable and clustered among higher-abundance cell types, obscuring their potentially powerful contributions to pathology or condition-specific variation. Finally, cells exist in a continuum of potentially overlapping identities, which are impossible to completely capture as distinct clusters—for instance, naïve, effector, and memory populations exist across CD4, CD8, and NK cell populations. Consequently, it is not possible to form accurate and discrete classes of cells. Instead, RISE offers an alternative perspective of cell “eigen-states”, which are multivariate and capture these distinct types of identities.

Both the strengths and limitations of RISE arise from the fact that it is a linear dimensionality reduction method. The method does not attempt to explicitly identify or correct for batch effects or other technical artifacts. Indeed, while there have been many attempts to understand and correct for these issues, there is a lack of agreement as to the extent of technical artifacts within scRNA-seq measurements, and many study designs do not enable a definitive line to be drawn between biological or technical variation^51,88^.

Instead, RISE can be used to identify patterns that are associated with batches; for example, by training a model on the condition factor matrix that predicts what batch a condition was processed in, we can understand which components reflect batch specific variation. By identifying component patterns, these gene expression alterations can be isolated and inspected. RISE requires the number of components as a model input; similarly to PCA, defining the optimal number of components can be difficult, with different heuristics being potentially appropriate depending on the goal of the modeling effort. In cases where one’s goal is to identify patterns predictive of disease, cross-validation using the factors as features may be appropriate. When there is not an association prediction or other task with which to evaluate the resulting factors, other metrics such as FMS or variance explained may aid in the task of selecting the size of the RISE model^37,89^.

An important strength of RISE is that it provides a foundational framework that can be expanded across various tasks. Additional constraints on the factors, such as nonnegativity, can enhance interpretation of the results^90^. The RISE algorithm can be altered to be a supervised factorization approach, leveraging information about each sample to better identify patterns that correspond to features on the cell or condition/sample scale^91^. RISE is well-suited to analyze single-omic analysis, but can be broadened to perform coupled decomposition for multi-omic data, such as combining transcriptomics with proteomics, to identify associative patterns across conditions^92^. With modest extension, such as constraints to account for spatial localization, RISE could be extended for identifying gene expression patterns in spatial measurements^49,93^.

In summary, we find that RISE is highly effective for modeling and exploring multi-condition single cell studies in both perturbational experiments and in an autoimmune cohort. RISE localizes variation to specific samples, genes, and populations of cells while enabling interpretation beyond what is possible with non-linear approaches such as autoencoders^14^. As the scale, capabilities, and questions asked using single cell techniques expand, fundamentally new approaches such as RISE that are scalable, easily interpretable, and unsupervised will be needed to understand how single cells coordinate their function within and across tissues^35^.

## Acknowledgements

This work was supported by NIH U19-AI172713 to A.S.M., an Emerging Leader Award from the Mark Foundation for Cancer Research, and an NSF Graduate Research Fellowship to A.R.

## Author Contributions

A.S.M. conceived of the study. A.R., B.O.-J., S.P., B.R., N.T., and A.S.M. performed the computational analysis. A.R., B.O.-J., S.P., B.R., N.T., and A.S.M helped to analyze the data. A.R., B.O.- J, S.P., A.A., and A.S.M. contributed to writing the paper.

## Supplemental Information

Supplemental Information is attached including plots for Figures S1–10 and descriptions for Figures S1–10 and Table S1–5. Content of Table S1–5 will be attached as Excel files.

## STAR Methods

### Resource Availability

#### Lead Contact

Further information and requests for resources should be directed to and will be fulfilled by the Lead Contact, Aaron Meyer.

#### Materials Availability

This study did not generate new unique reagents.

#### Data and Code Availability

- All analysis was implemented in Python v3.12 and can be found at https://github.com/meyer-lab/scCP, release 1.0.
- The implementation of RISE is available at https://github.com/meyer-lab/parafac2. The single-cell gene expression was obtained from Figshare (https://doi.org/10.6084/m9.figshare.11837097) and via GEO accession number GSE174188.
- Any additional information required to reproduce this work is available from the Lead Contact.

## Method Details

### PARAFAC, PARFAC2, and RISE

Our implementation and interpretation framework around the PARAFAC2 tensor decomposition algorithm, which has been optimized for its use towards scRNA-seq, is coined Reduction and Insight in Single-cell Exploration (RISE). We use RISE to refer to our adaptation of PARAFAC2 which includes parameter initialization scheme for PARAFAC2, parameter selection of PARAFAC2, use of Nesterov acceleration for optimization, and interpretation utilities of the PARAFAC2 factors.

#### PARAFAC

As PARAFAC and PARAFAC2 serve as the core modeling approach behind RISE, we will first provide a mathematical explanation of PARAFAC. PARAFAC is a generalization of principal components analysis, which primarily operates on second-order tensors (matrices). Despite its applicability to arbitrary-order tensors, we will explain the usage of PARAFAC to third-order tensors, as this is most relevant to our subsequent PARAFAC2 explanation. Given a third-order tensor, ***X***, of shape *I* × *J* × *K*, PARAFAC seeks to approximate ***X*** as a sum of component rank-one tensors. This set of component rank-one tensors are collectively represented in factor matrices: *A* of shape *I* × *R, B* of shape *J* × *R*, and *C* of shape *K* × *R*, where *R* is the number of components into which ***X*** is decomposed. Specifically, PARAFAC seeks

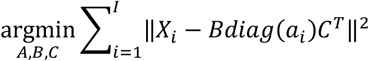

where *X*_*i*_ is the *i*^th^ slice of ***X*** along the first mode, and *diag*(*a*_*i*_) is the diagonal matrix whose nonzero entries are equal to the *i*^th^ row of *A*.

#### PARAFAC2

In single-cell transcriptomics, a different set of cells is profiled for each condition, and the number of cells profiled in each condition may vary. This precludes the direct arrangement of these measurements into a single tensor, and prevents the use of PARAFAC and many other existing tensor-based decomposition methods^94^. This data would instead be better represented as a length-*I* list, ***X***_***sc***_, of second-order tensors, 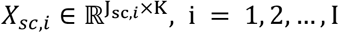, where *I* is the number of conditions, *J*_*SC,i*_ is the number of cells profiled in condition *i*, and *K* is the number of genes. PARAFAC2 explicitly handles data of this form, where one axis (in this case the cell axis) varies across another (here, the condition axis). We use the direct fitting algorithm originally proposed by Kiers et al. to fit the PARAFAC2 model^37^.

In fitting the PARAFAC2 model, we seek

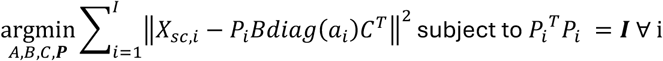

where 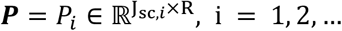, *I* are the projection matrices, where *R* is the rank of the PARAFAC2 decomposition; *diag*(*a*_*i*_) is the diagonal matrix whose nonzero entries are equal to the *i*^th^ row of *A* ∈ ℝ^*I*×*R*^, the condition factor matrix; *B* ∈ ℝ^*R*×*R*^ is the cell eigen-state factor matrix; *C* ∈ ℝ^*k*×*R*^ is the gene factor matrix; and ***I*** is the identity matrix^37^.

Comparing the objective functions for PARAFAC and PARAFAC2, one can see that the difference between the two models is simply the addition of the orthonormal projection matrices ***P*** in PARAFAC2, which accounts for data that is unaligned in one dimension, such as scRNA-seq.

We fit PARAFAC2 using alternating least-squares, whereby ***P*** and the factor matrices, *A, B*, and *C*, are alternately fit. First, the factor matrices are updated as

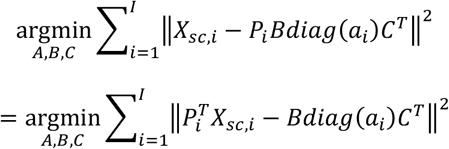

which is equivalent to running PARAFAC^94^ on the single, aligned, third-order tensor whose *i*^th^ slice along the first mode is equal to 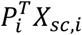. Then, the projection matrices, *P*, are updated by solving the Orthogonal Procrustes problem. Given matrices *X*, Y ∈ ℝ^n×m^, the Orthogonal Procrustes problem seeks an orthogonal matrix *Q* ∈ ℝ^n×n^ that minimizes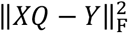. The solution is given by *Q* = UV^T^, where UΣV^T^is the singular value decomposition of Y^T^*X*. For a low-rank decomposition, we update each ***P***_***i***_ ∈ ***P*** as the product of the first *R* left and right singular vectors from the SVD of 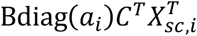. This process of updating the factor matrices using PARAFAC and the projections via Orthogonal Procrustes is repeated until a specific number of iterations elapses or the difference in R2X between iterations reaches a threshold.

#### RISE

##### Parameter Initialization

Parameter initialization was performed by using the SVD of the matrix-flattened data, as previously described^37^. Each of *X*_*sC,i*_ was concatenated along the cell dimension to form a cell by gene matrix containing cells from every condition. Randomized SVD^95^ was performed on this matrix and the first *R* right singular vectors were used to initialize *B. A* was filled with ones and *C* was initialized to the identity matrix.

##### Optimization Technique

We used the TensorLy implementation of PARAFAC and alternating least squares (ALS) to solve for each factor, wherein each mode is iteratively solved using least squares^96^. We used 20 iterations of PARAFAC per PARAFAC2 iteration. We terminated the PARAFAC2 fitting algorithm once either 200 iterations had elapsed or the difference in R2X between iterations was less than 10^−6^.

To speed the fitting process, we use all-at-once Nesterov-like acceleration for fitting the model as previously described^97^, with extrapolation step size determination parameters of 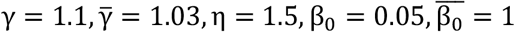.

##### Post-Factorization Alignment

Like PCA, CP-based decomposition component patterns are made up of the product of vector weights for each dimension in the data. CP-based decompositions generally provide a unique solution, aside from three indeterminacies: the sign of the factors, scaling between factors, and ordering of the components. We therefore applied rules to yield consistent factorization results, helping with the interpretation of the results. To address the scaling indeterminacy, we normalized each factor matrix to have components of unit scale. The sign indeterminacy was partially resolved by making the conditions factors positive.

PARAFAC2 factors are order indeterminant, where the order of each component can be arbitrarily arranged. ***A***, the conditions factor, was used to arrange components according to their Gini coefficient of association across the conditions, so that the first components were generally found to be associated with all the conditions, while the later components were generally condition specific.

### Reconstruction Error

We used the metric reconstruction error (R2X), or the percentage of variance explained by the model, to assess the model fit. First, the Frobenius norm squared of the difference is calculated between the data (*X*_*data*_) and the reconstruction based of the factors and projections (*X*_*Pf*2_). This difference is normalized by the Frobenius norm of the data and subtracted from 1 to result in a range from 0 to 1, where 1 represents 100% variance explained.

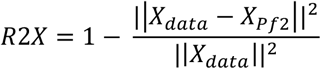

### Factor Match Score

The factor match score (FMS) measures the similarity between two sets of factors^49^. Two FMS strategies were implemented. First, to determine how the size of the data influences the factors, the FMS was compared between RISE factorizations with the same number of components on the entire dataset and with various fractions of the cells removed. Second, to determine the stability of the factorization at varying numbers of components, cells were bootstrapped to create a dataset of the same size. The FMS was then calculated for the original dataset and the reconstructed dataset at various components. Thus, after 2 unique RISE factorizations are applied in both cases, the FMS compares the conditions (*A*) and genes (*B*) factor matrix of the original factorization to the reconstructed factor matrices, 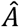 and 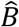. The eigen-states factors vary in what cells they represent, so they are not included in calculating the FMS. An FMS of 1 indicates complete similarity. A linear sum assignment is also applied because a decomposition can have the same components, but in a different order.

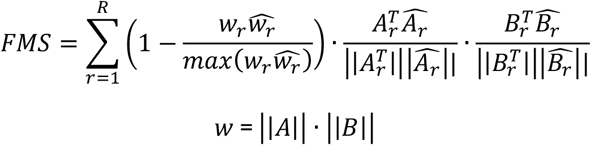

### Cell Type Abundance Change Simulations

To stimulate experiments where a cell type is not present, we removed cells labeled as a specific cell type and determined if RISE could identify this change. First, RISE was applied to all the cells and conditions of a dataset. Then, the cells of X cell type were removed for a specific experiment, Y, and RISE was applied to this cell type-perturbed dataset. After comparing the canonical genes highly weighted for the gene factors for both analyses, the conditions’ component corresponding to canonical markers of the cell type were identified and compared between the two cases. To stimulate an experiment with fewer cells of a certain cell type than other conditions, we subsampled the given cell type by the indicated amount, rather than removing all cells.

### Data Normalization, Processing, Cell Annotation, and Visualization

#### Perturbation Study Analysis

Data Normalization and Processing: The single-cell gene expression was obtained from Chen et al^15^. The first step before normalization was removing doublets with the doubletdetection package applied to the raw RNA counts matrix^98^. Cells with a doublet probability of 0.5 or more were removed from the dataset.

Once doublets were removed, the data was normalized. Genes with less than 0.01 mean raw expression across all cells were removed. Each cell was then normalized by read depth using CPM normalization. Next, each gene was scaled according to its sum. Finally, the data was log transformed via the Delta method,

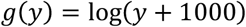

where g(y) is the transformed gene and y is the pre-transformed gene^99^. Finally, each gene was mean-centered. The final size of the dataset included 29433 cells and 9461 genes. The data was then fit using the RISE model with 20 components as described above. PaCMAP was used on the concatenated matrix of the projections across all conditions to visualize the cell-to-cell variation and embed cells into low-dimensional space, using default parameters of n_neighbors=10, MN=0.5, and FP_ratio=2.0^50^.

Cell Type Annotation: Lower-resolution cellular clusters were defined by Leiden clustering of the RISE projections, resulting in 6 Leiden clusters, with parameters n_neighbors=10 for the nearest neighbors distance and resolution=2.5 for the Leiden algorithm^100^. To provide each cluster with a cell type annotation, genes expressed uniquely within each cluster were found using differential expression analysis by the dispersion-based method with parameters of min_mean=0.005, max_mean=10, min_disp=0.5^101^. These genes were used to annotate each cluster based on their overlap with previously reported marker genes^15^. After the lower-resolution assignments, a finer-grained annotation was performed based on each cluster’s expression of a new set of marker genes (Table S5).

#### SLE Analysis

Data Normalization and Processing: Previous analysis used batch corrected data^62^. To prevent potential sources of bias, we used the raw RNA counts matrix after doublet analysis by the original authors. The same normalization pipeline was used as above, except genes with less than 0.1 mean raw expression across all cells were removed. The final size of the dataset was 1,263,673 cells across all conditions and 2,161 genes. The data was then fit using the RISE model with 30 components as described above.

Cell Type Annotation: Once again, a similar process was used to assign lower and higher resolution cell types as above. Nearest neighbors distance with n_neighbors=10 and resolution=3 for the Leiden algorithm was used to detect 47 Leiden populations. Low and high-resolution cell types were assigned based on a set of marker genes included in Perez et al (Table S5)^62^.

### Logistic Regression

Scikit learn was used to train an L1-penalized logistic regression model to predict SLE status using the RISE condition factor matrix. A nested cross-validation strategy was initially copied from Perez et al., where the model was trained on the samples of one processing batch and then predicted the rest of the samples in each of the three remaining batches^102^. In addition, an alternative, more traditional nested cross-validation strategy was performed that trained three processing batches and predicted the fourth batch. After the batches are split based on the cross-validation strategy, sample-wise 5-fold inner cross-validation was performed to optimize the L1-penalty of the classifier. Solving was performed with the Stochastic Average Gradient (SAGA) solver, a tolerance of 1 × 10^−4^, and a maximum iteration of 10,000^103^. Both the prediction accuracy and ROC AUC were used as a metric to measure the ability to associate the factors with SLE status. Regularization strength was fit by grid search using the cross-validation implementation described above.

### Cell Type Composition and Pseudobulking

Cell type composition was analyzed by calculating the percentage of all cells in one experimental condition that corresponded to that cell type. Pseudobulk profiles were created by taking the average of the normalized expression for each cell type in each experimental condition.

### Principal Components Analysis

Principal components analysis (PCA) was performed using scikit learn^102^. The data was “flattened” into a two-dimensional matrix of all cells by genes. The number of components was set to match RISE.

### Gene Set Enrichment Analysis

The 30 most positively and negatively weighted genes for each component were submitted to ToppGene. ToppGene applies a fuzzy-based similarity measure to calculate semantic similarities between genes. To determine gene annotation significance, p-values were generated using random sampling across the entire genome, based on the empirical distribution of the similarity scores. Significantly (FDR < 0.05) enriched biological processes were retained^104^.

