## Supplement for "Integrative, high-resolution analysis of single cell gene expression across experimental conditions with PARAFAC2-RISE"

**Table S1: Comprehensive summary of traditional single-cell analysis techniques and how it differs from RISE.**

**Table S2. RISE components associate each pattern to specific genes for the scRNA-seq perturbation dataset.** (a-b) The top 30 most (a) negatively and (b) positively weighted genes identified by RISE for all components, for analyzing the drug perturbation dataset with a 20 component RISE model.

**Table S3. Overview of the RISE component interpretation for the scRNA-seq perturbation dataset.** Description of each component for the cell types that are highly weighted for the weighted projections and the top five highly weighted genes, along with the sign for each case. The gene set enrichment analysis (GSEA) biological process results for each component gene module (top 30 weighted genes) were noted alongside the correspondence to a cell type.

**Table S4. RISE components associate each pattern to specific genes for scRNA-seq SLE and healthy cohort studies.** (a-b) The top 30 most (a) negatively and (b) positively weighted genes identified by RISE for all components, for analyzing the SLE dataset with a 30 component RISE model.

**Table S5. Canonical gene markers used to assign Leiden clusters to immune cell types for the scRNA-seq perturbational and SLE datasets.**

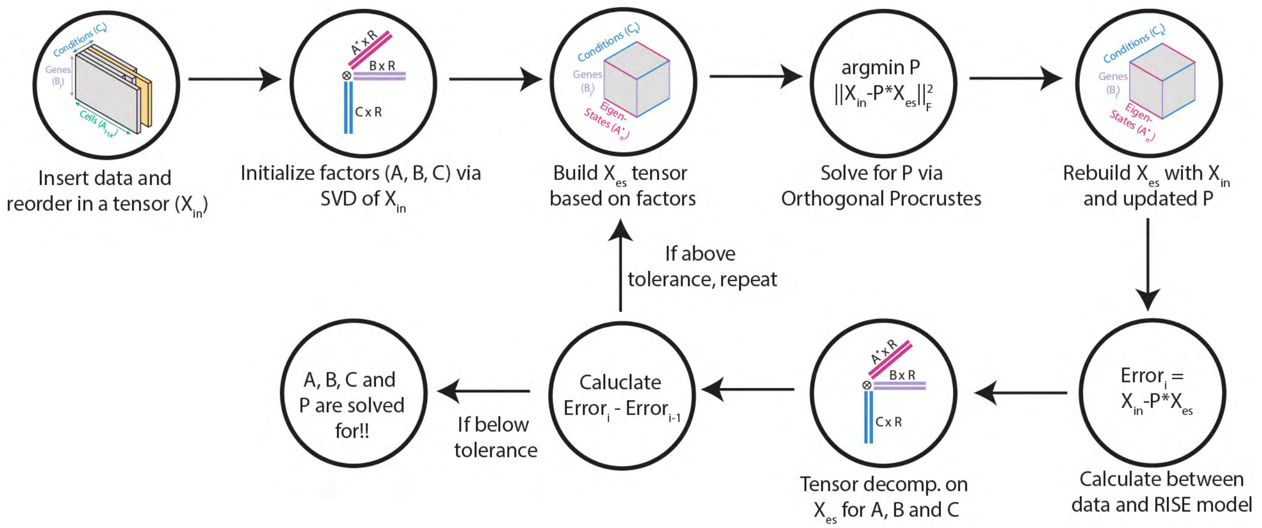


**Figure S1. The RISE algorithm iterates between solving for the factors and projections until convergence.** The algorithm optimizes the error between the factors (A, B, C) for the conditions, eigen-states and genes with PARAFAC, and then the data is re-projected onto the common reference frame. This process is repeated until convergence.

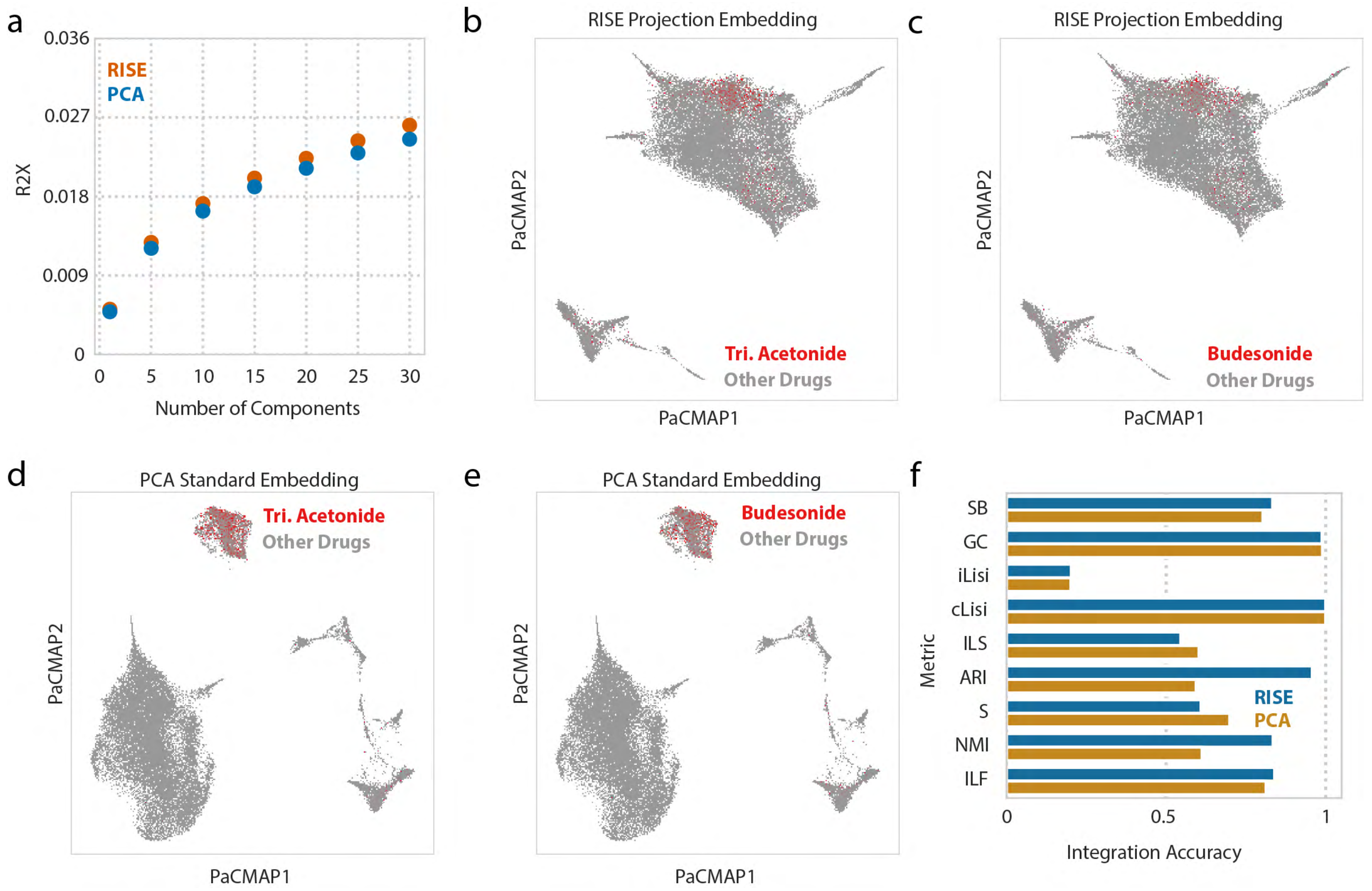


**Figure S2. RISE prevents condition-specific embedding clusters. (a)** Fraction of the variance explained by PCA and RISE for each number of components. **(b–e)** PaCMAP embedding of the (b–c) RISE projections and (d–e) PCA embedding on the flattened data, with cells from the (b, d) triamcinolone acetonide and (c, e) budesonide treatment condition highlighted. **(f)** Integration accuracy for batch integration and biological conservation for PCA and RISE embedding; batch integration metrics include k-nearest-neighbor graph connectivity (GC), batch average silhouette width (SB), and graph integration local inverse Simpson's Index (iLisi); biological conservation metrics include adjusted rand index (ARI), normalized mutual information (NMI), cell-type average silhouette width (S), isolated label silhouette (ILS), cell type local inverse Simpson's Index (cLisi), and isolated label F1 (ILF).

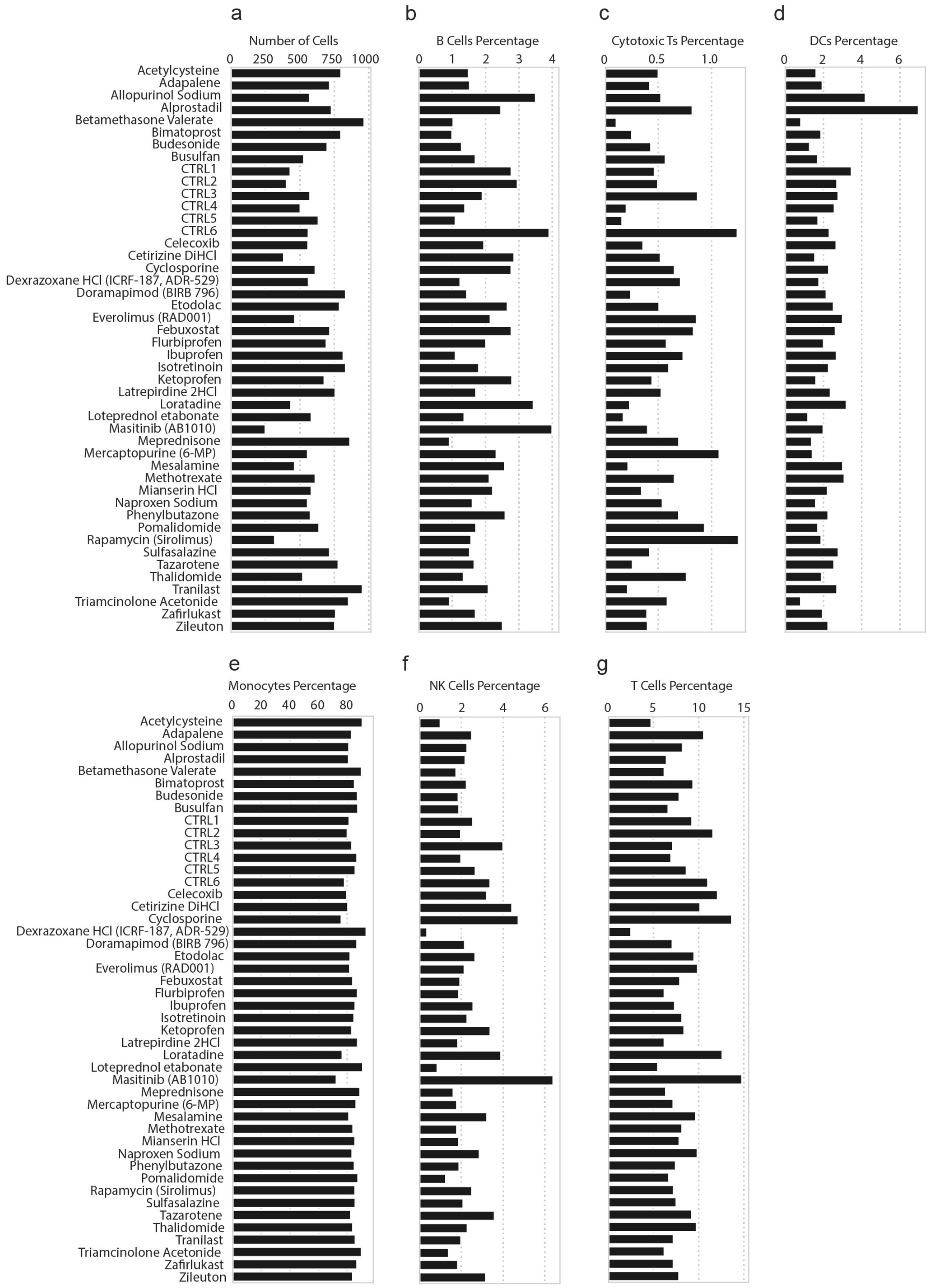


**Figure S3. RISE embedding and post-clustering results in similar percentages of low-resolution immune cell types across all conditions. (a)** Number of cells across all drug perturbations and controls. **(b-g)** Cell type percentage across all drug perturbations and controls, broken down by low resolution immune cell types, including (b) B cells, (c) cytotoxic T cells, (d) DCs, (e) monocytes, (f) NK cells, and (g) T cells.

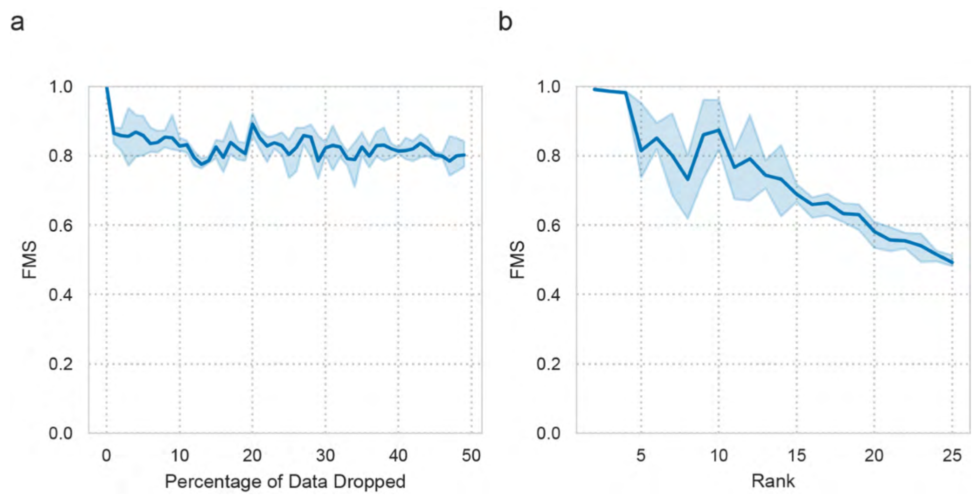


**Figure S4. RISE results in consistent factor patterns despite reduced numbers of cells or varying component numbers.** **(a)** Factor match score (FMS) when comparing the RISE gene and condition factors of the full drug perturbational scRNA-seq dataset or the same dataset with a reduced subset of cells. A rank of 20 of was used for both datasets. **(b)** FMS comparing the full dataset to a bootstrapped dataset of the same size, at various ranks.

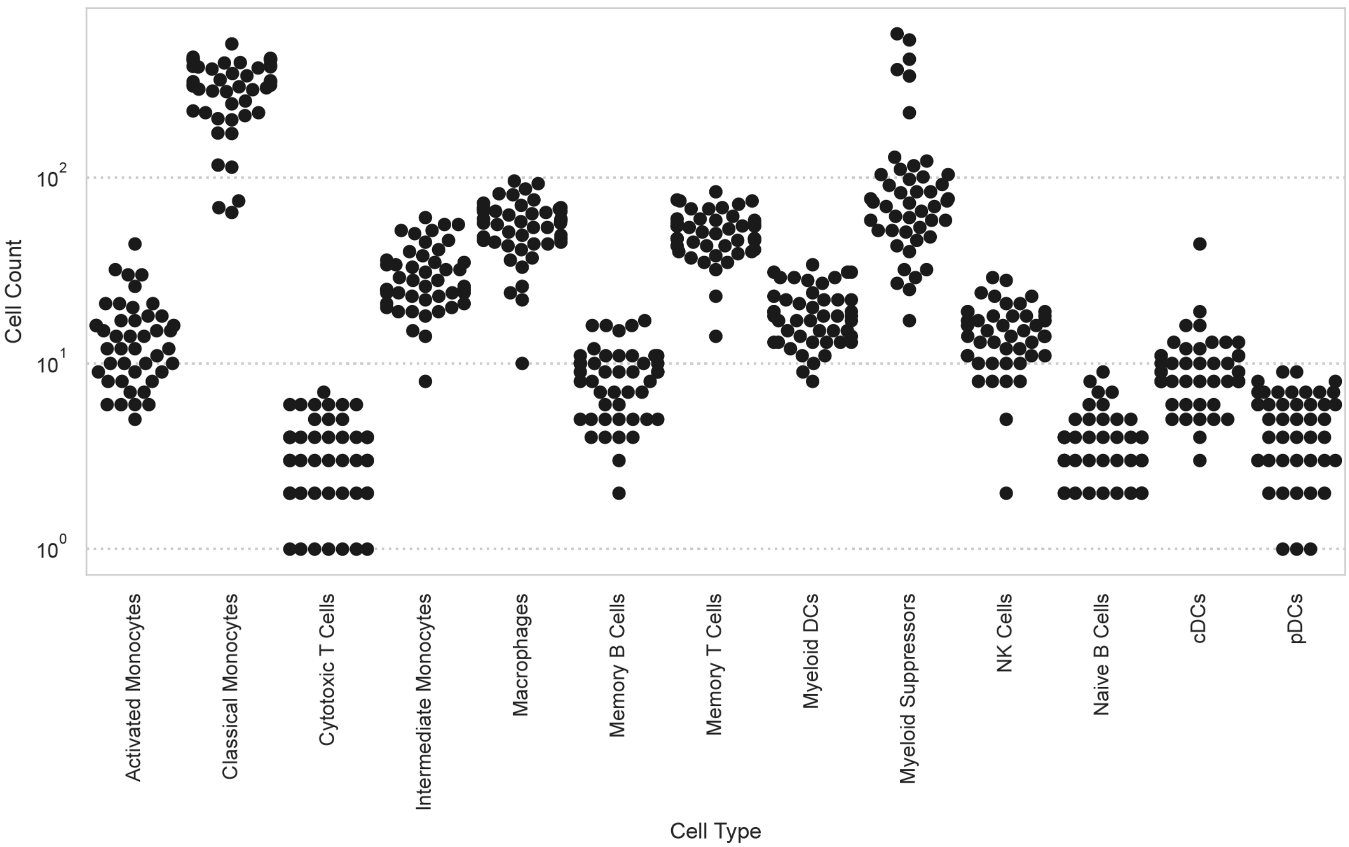


**Figure S5. RISE embedding and post-clustering results in similar abundances for the high-resolution immune cell types across conditions. (a-m)** Cell counts across each drug perturbation and control, stratified by high-resolution immune cell type.

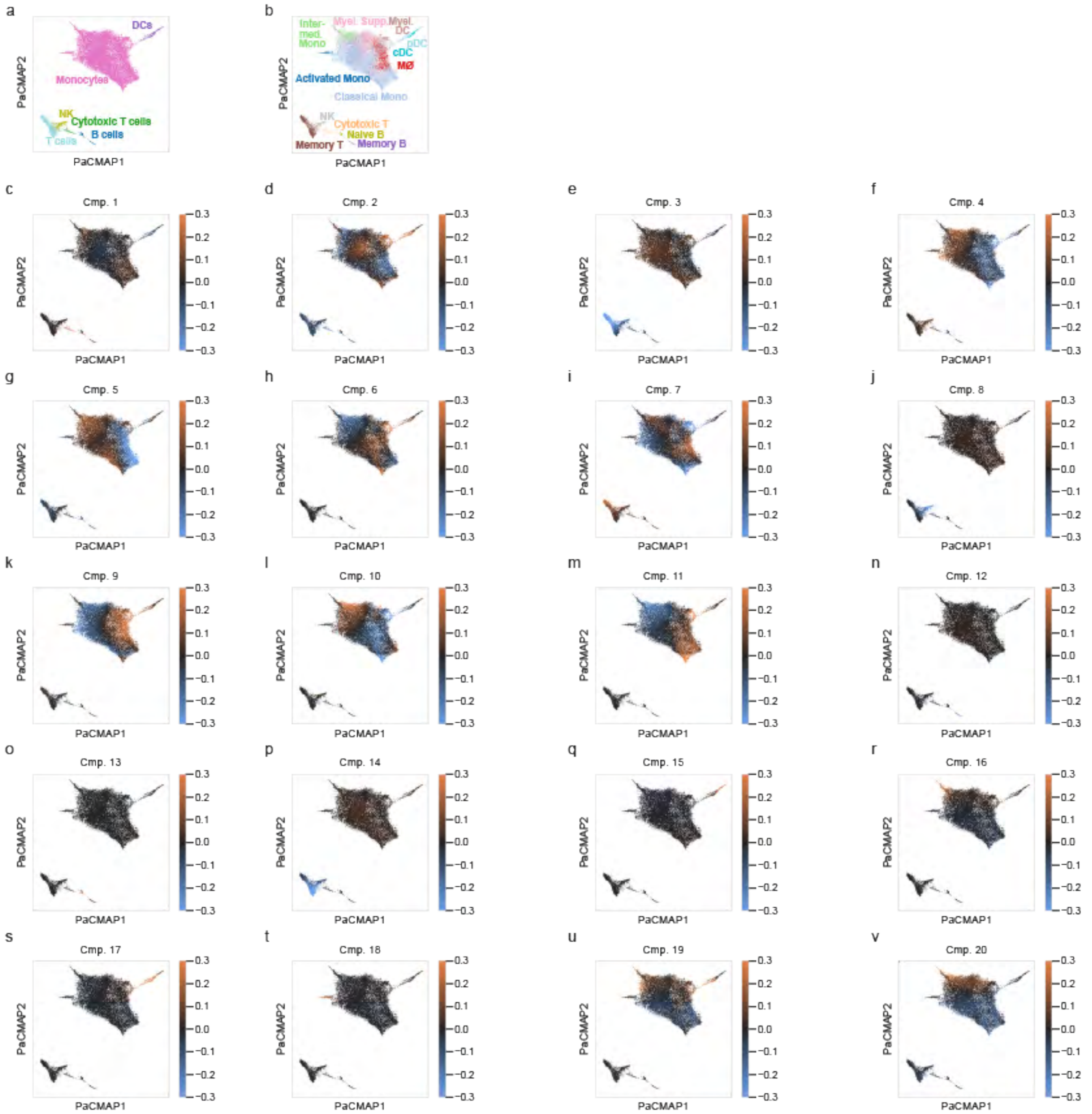


**Figure S6. RISE cell weights associate each component to specific cell populations. (a-b)** PaCMAP of the RISE projections, with cells colored by their (a) low-resolution and (b) high-resolution cell type annotations. **(c-v)** PaCMAP of the RISE projections, with cells colored by the weighted projection of each component, for the drug perturbation dataset with a 20 component RISE model.

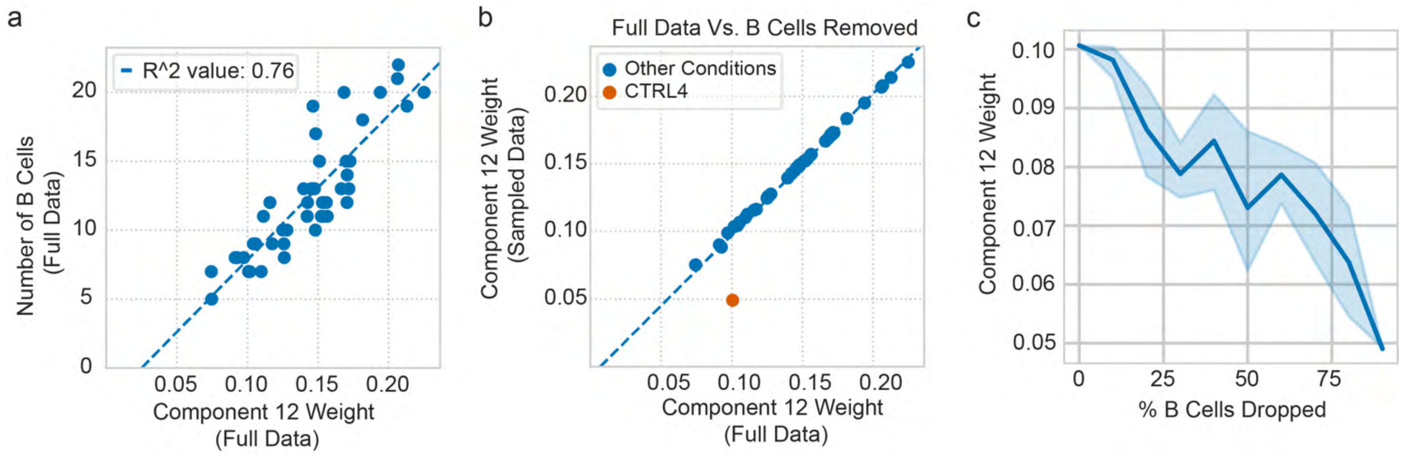


**Figure S7. RISE components represent specific subsets of cells with unique gene expression responses. (a)** Relationship between B cell count and component 12 weight for each experimental condition. **(b)** Comparison of the component 12 condition weights when RISE is applied to the full dataset to one in which B cells were removed from scRNA-seq condition CTRL4. **(c)** RISE component 12 condition weights when varying fractions of B cells are removed.

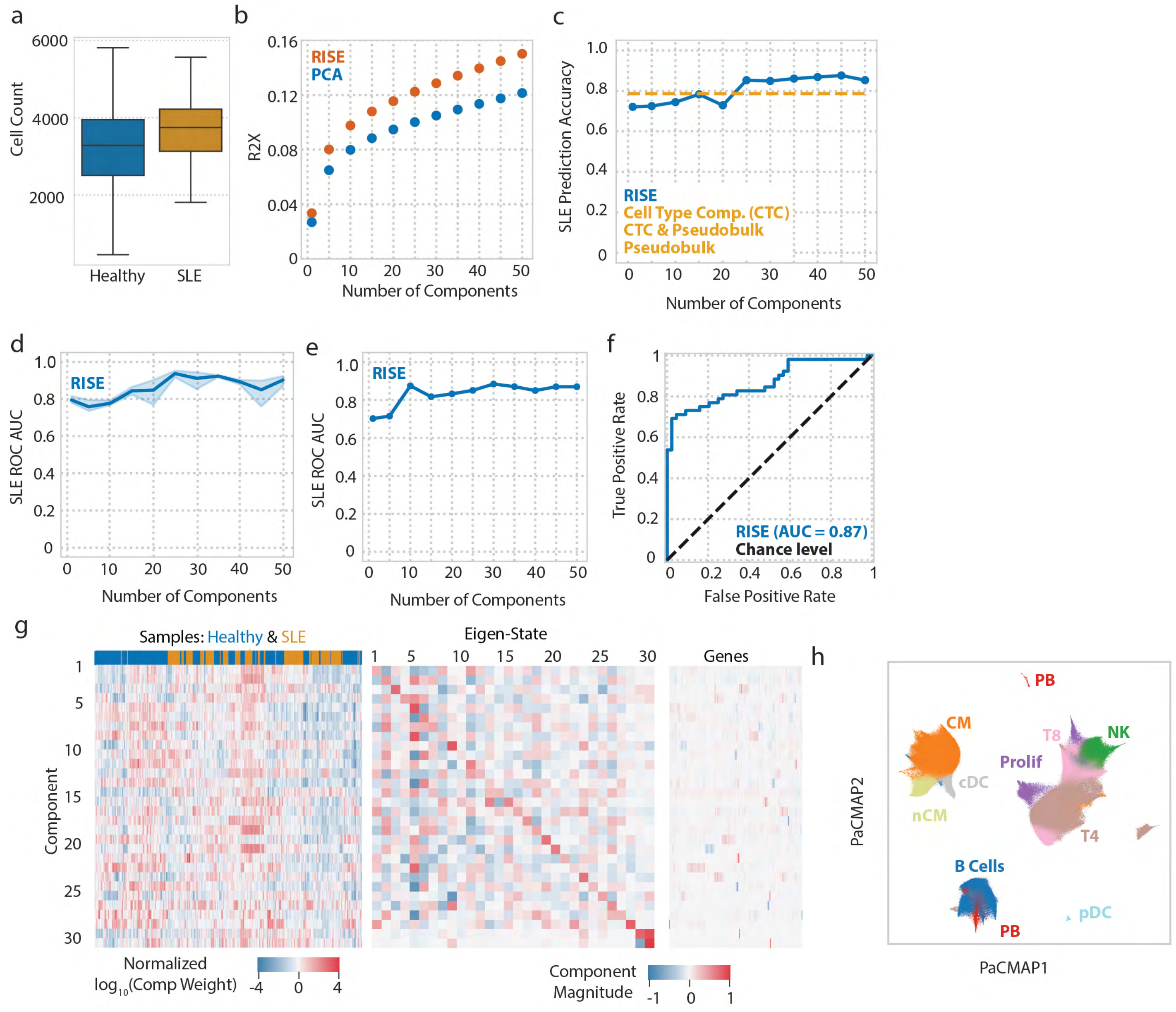


**Figure S8. RISE identifies patterns associated with patient status and separates immune cell populations. (a)** Total cell count per experimental sample for SLE and healthy samples. **(b)** Percent variance explained using different numbers of components with PCA versus RISE. **(c)** Accuracy of predicting SLE status using the condition factors as the input for logistic regression with a L1 penalty across different numbers of components. **(d)** ROC AUC of a logistic regression classifier, using bootstrapped patient factors, predicting SLE status. **(e)** ROC AUC of a logistic regression classifier, using the patient factors and an alternative cross-validation strategy, predicting SLE status. **(f)** ROC curve of a logistic regression classifier, using an alternative cross-validation strategy, predicting SLE status with a 30-component RISE model. **(g)** The (left) samples, (middle) eigen-state, and (right) gene factors for each component. The genes factor matrix is filtered for genes having an effect magnitude of 0.8 or more within one of the components. The sample factors are non-negative and have been log transformed to aid comparisons. **(h)** PaCMAP of the RISE projections, with cells colored by the low-resolution cell type annotations.

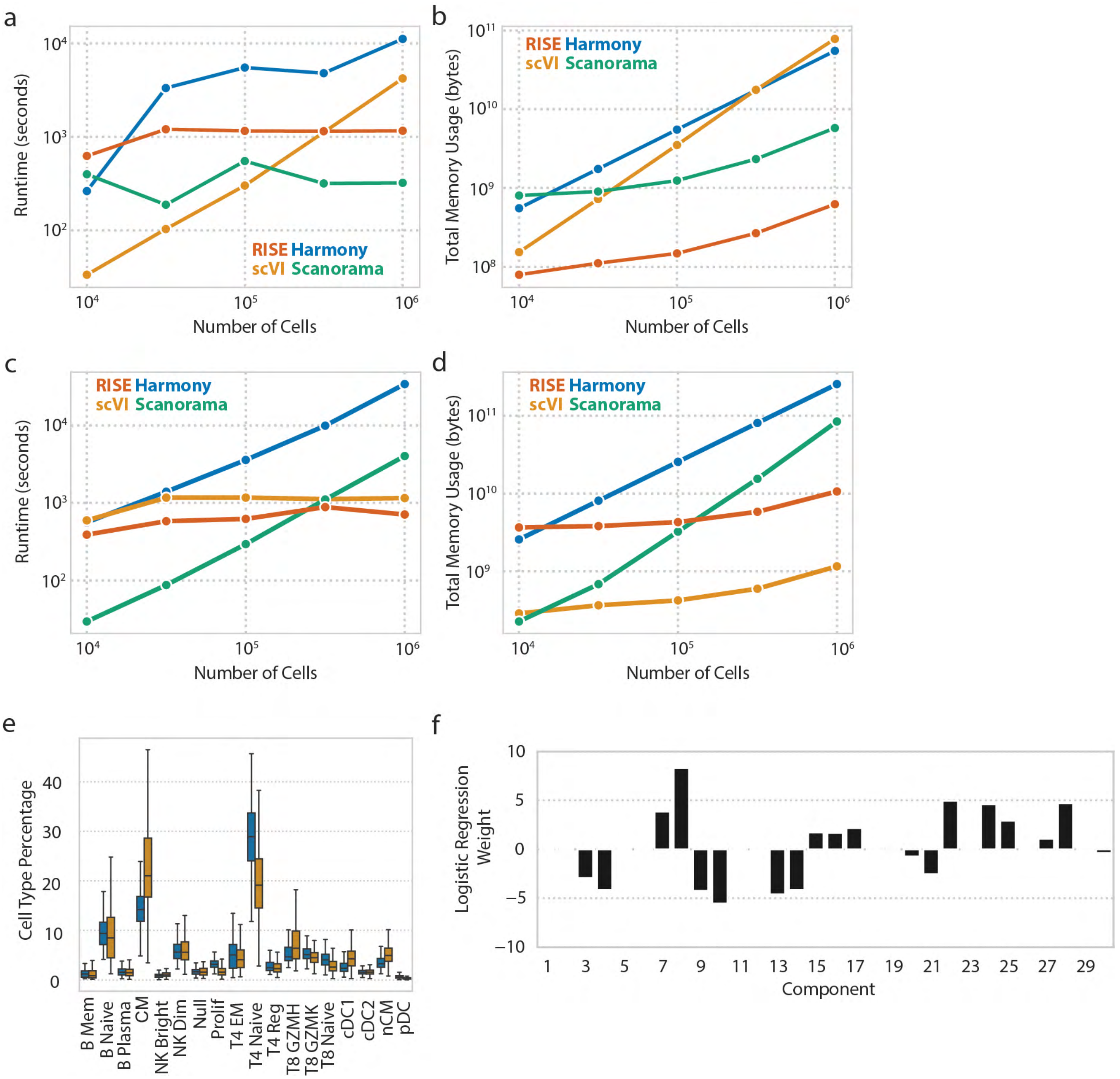


**Figure S9. RISE's runtime and total memory usage across different number of cells and genes are comparable to other methods and RISE identifies components associated with lupus status. (a, c)** Runtime comparison in seconds of RISE, scVI, Scanorama, and Harmony across different numbers of cells with ~2000 genes. **(b, d)** Total memory usage comparison in bytes of RISE, scVI, Scanorama, and Harmony across different numbers of cells with ~10,000 genes. Total memory usage is the sum of CPU and GPU memory usage. This analysis was conducted in Python, and the default parameters for each method's Python package implementations were used. **(e)** Abundance of the high-resolution cell type annotations across SLE and healthy samples. **(f)** Weights of the logistic regression model (L1 penalty) predicting SLE status using the samples factor matrix.

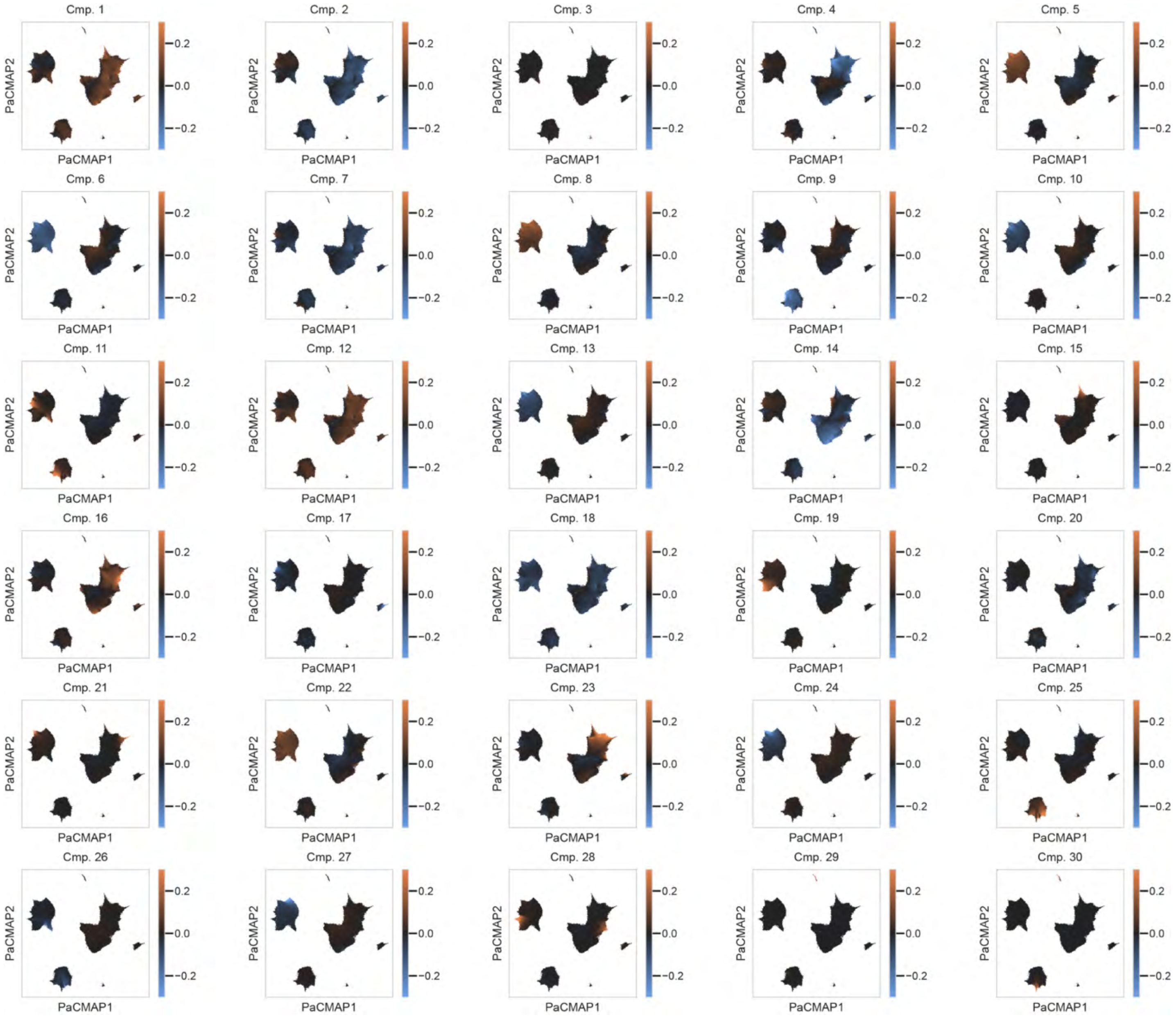


**Figure S10. RISE components associate each pattern to specific cell populations.** PaCMAP of the RISE projections, with cells colored by the weighted projections for the indicated component. Analysis of the SLE dataset with a 30 component RISE model.
